# Agent-based modelling of a nematode system provides novel insights into the evolutionary constraints and modulators of phenotypic plasticity, bet-hedging, and environmental homeostasis

**DOI:** 10.64898/2026.06.02.729670

**Authors:** Riccardo Tarantino

## Abstract

In this study, I implemented an agent-based model aimed at exploring the competition between plastic and non-plastic genotypes of the dimorphic nematode *Pristionchus pacificus* in a digital environment with periodic fluctuations in food resources. Simulation scenarios monitoring frequency and time until fixation of two alleles of the developmental switch gene *eud-1* were performed to capture emerging eco-evolutionary patterns generated by the interplay of three main variables, namely the intrinsic cost of plasticity, the timescale of environmental fluctuations, and the individual degree of plasticity. Interestingly, while intermediate-to-long periods of environmental stability and higher levels of plasticity might favour plastic strategists in a cost-free condition, the introduction and increase of inherent costs of plasticity could select for genotypes with either very low or high sensitivity to environmental cues, induce a sequential collapse in the frequency of fixation of plastic strains and time of coexistence between strains, and make invasions by non-plastic mutants/immigrants more likely until a plateau is reached. In addition, asymmetries in fitness between the two alternative phenotypes might be an almost necessary condition to enable the invasion of a non-plastic population by plastic genotypes. Collectively, while confirming previous theoretical findings, these outcomes may also uncover the sensitivity of a nematode system involving stochastic, conditional, and constitutive phenotype production to even small changes in key variables, suggesting the existence of phase transitions and critical evolutionary constraints.

## 1. Introduction

The relationship between genotype and phenotype is one of the cornerstones of biology, and the environment often plays a pivotal role in directing the process of conversion of genetic information into traits at every phenotypic level, from proteins up to complex behaviours. Indeed, traits may either remain stable or be expressed differently in response to environmental stimuli. The former property is known as *environmental homeostasis* (Pigliucci, 2010), while the latter is called *phenotypic plasticity*, that is commonly defined as the ability of a genotype to produce different phenotypes in response to different environmental conditions and is an important, astonishingly widespread strategy which may enable survival in rapidly changing ecological contexts or allow the colonisation of new niches (Pigliucci, 2001; Ghalambor et al., 2007; Fusco & Minelli, 2010; Bonamour et al., 2019). The pervasiveness of plasticity both among prokaryotes and eukaryotes is clear when looking at the impressive number of case studies reported in the literature and referring to a broad variety of taxa, from single-celled organisms to trees or mammals. Remarkable examples of plasticity include regulation of molecule production in bacteria in response to cell density (Kummerli et al., 2009), repeated modifications of body length in marine iguanas depending on food availability (Wikelski & Thom, 2000), development of a different number of segments in centipedes based on the temperature experienced during embryogenesis (Vedel et al., 2008), and countless others involving the most diverse traits, either with continuous or discrete variation (*polyphenism*), triggering factors, and potential for reversibility of the response (Bento et al., 2010; Lin et al., 2018; Escobar-Sandoval et al., 2021; Mu et al., 2021; Sjulgård et al., 2021; Fogg et al., 2022; Marian et al., 2024).

Interestingly, while plasticity is typically regarded as a conditionally regulated property, some authors include stochastic regulation as a relevant driver of plastic responses in both unicellular and multicellular organisms, as shown by species exhibiting both regulations simultaneously (Sommer R. J., 2020). Indeed, environment-dependent plasticity can evolve in combination with *bet-hedging*, that is, a risk-spreading strategy – promoted by unpredictable environments and between-generations variation (Donaldson-Matasci et al., 2013) – which leads to a reduction in mean fitness while offering a beneficial trade-off across generations by reducing fitness variance (Yasui & Garcia-Gonzalez, 2016; Susoy & Sommer, 2016). An example of this mixed strategy, also known as coin-flipping plasticity, was reported in pea aphids (Grantham et al., 2016): although the probability to produce winged or wingless offspring in this species depends on the level of crowding experienced by asexual females, both phenotypes are produced by individual aphids. This case study effectively illustrates the complexity of plasticity and its regulation within biological systems.

Most importantly, while metabolic, physiological, and developmental processes are intrinsically sensitive to some environmental variables (Nijhout, 2002), the genetic architecture and processes underlying plasticity – including transcriptional, post-transcriptional, and epigenetic regulation – provide the substrate for the evolutionary forces to act upon. (Chevin et al., 2013; Peltier et al., 2018; Fortuna, 2022; Kovuri et al., 2023). For instance, the magnitude of plastic responses across environmental conditions (i.e., the *degree of plasticity*) is altered by single-nucleotide mutations in yeasts (Duveau et al., 2017) and nematodes (Vigne et al., 2021); furthermore, this trait has been shown to be modulated by selection in both laboratory settings (Walworth et al., 2016; Mallard et al., 2020) and wild populations (Nussey et al., 2005). The latter case studies clearly demonstrate that evolutionary processes can indeed regulate plasticity, leading to an increase, maintenance, or reduction of its level, including the complete loss of this ability.

These dynamics may be governed by the impact of trait plasticity on fitness. More specifically, plasticity may be maladaptive, neutral, or adaptive (Ghalambor et al., 2007), depending on whether it has negative, null, or positive effects on fitness, respectively. For instance, it may be favoured by natural selection when organisms face heterogeneous or variable environments, assuming that i) the plastic response moves the mean phenotype closer to the phenotypic optimum of each environment, ii) environmental cues are sufficiently reliable, and iii) the *cost of plasticity* is low. Particularly, while often challenging to detect, such cost – which might be related to additional regulatory machinery (Murren et al., 2015; Hendry, 2015) – is a potentially decisive factor affecting evolutionary outcomes. In these conditions, plasticity can be considered a generalist strategy (Chevin et al., 2013). In contrast, environments characterised by infrequent variation or unpredictable patterns of variation select for fixed, unconditionally expressed traits. As long as it confers the highest fitness in a specific environment, constitutive expression of a trait – which might evolve from a reduction or loss of plasticity if the environmentally induced response moves the mean phenotype further from the optimum – is considered a specialist strategy (Acasuso-Rivero et al., 2019; Storz & Scott, 2021). Finally, plasticity may have no effects on fitness, so its evolution is likely to depend on stochastic mechanisms such as genetic drift (Gomulkiewicz & Stinchcombe, 2022).

Along with research using experimental evolution (reviewed in Kassen (2002)) and analytical treatments (Gabriel & Lynch, 1992; van Tienderen, 1997; King & Hadfield, 2019; Kasada & Yoshida, 2020), computational modelling and simulations constitute an important approach that has shed light on the conditions driving the competition between plastic and non-plastic genotypes. For instance, the potential of digital organisms for testing theoretical predictions including the hypothesis that variable and static environments select for and against plasticity, respectively, has recently been highlighted (Fortuna, 2022). Using the Avida Digital Evolution Platform, Lalejini and Ofria (2016) investigated the selective pressures leading to plasticity and, in addition to obtaining outcomes consistent with previous assessments including Ghalambor et al. (2010), they also observed that traits were generally expressed unconditionally prior to the evolution of conditional trait expression and reported the evolution of sub-optimal forms of plasticity before optimal forms. Other insights of general interest can be found in another study involving digital plastic and non-plastic organisms facing different environmental conditions (Miras, 2024), i.e., that genetic costs of plasticity might undermine the potential benefits of this strategy. A similar approach is also found in research using *agent-based modelling*. For instance, Edelaar et al. (2017) explored the conditions leading to either adaptive plasticity or matching habitat choice as possible solutions to environmental heterogeneity with the simulation platform NetLogo, while Kalirad & Sommer (2024) embraced an integrative theoretical-experimental framework by combining this modelling technique and laboratory measurements taken from the nematode species *Pristionchus pacificus* to investigate the effects of plasticity and stochasticity on competition and coexistence of different strains.

Agent-based modelling is a technique used to implement dynamic computational environments (i.e., agent-based models, hereafter referred to as ABMs) where the global behaviour of complex systems can be simulated as a product of multiple interactions among independent, proactive individuals called *agents*. In a nutshell, the modeller assigns agents specific properties, state variables, and algorithms, and the subsequent execution of the program determines all the activities the agents will perform within their virtual environment. Most importantly, the simultaneous performance of these tasks can lead to the spontaneous formation of emerging patterns of potential interest (for further details on ABMs and complex systems, see Wilensky & Rand (2015)). The interplay between agents and environments makes this kind of modelling particularly suitable for applications in ecology and evolution (Murphy et al., 2020), where changes in the gene pool of a population may be seen as a collective phenomenon produced by multiple generations of organisms following a set of hereditary and developmental rules in different ecological contexts. More specifically, ABMs may be appropriate for drawing general insights into the selective pressures favouring plasticity, stochasticity, or environmental homeostasis by simulating hypothetical scenarios involving agents characterised as their real counterparts.

In line with the integrative approach adopted by Kalirad & Sommer (2024), I implemented an ABM to carry out an exploratory analysis aimed at investigating competition between plastic and non-plastic genotypes using a virtual environment populated by agents whose behaviours and state variables are grounded in the experimental data available for the model organism *Pristionchus pacificus*. Despite the limitations related to the specific characteristics of the chosen organism, this focus allows the inclusion of an additional dimension into the study, namely a stochastic component involved in phenotype production — a phenomenon that has been extensively documented in this species (Susoy & Sommer, 2016) — which provides material for an assessment of the interplay between responses induced by the environment and bet-hedging. In this study, I was interested in exploring: i) the specific timescales of environmental fluctuation and the genetic conditions increasing the chance for the plastic genotype to become prevalent, ii) the sensitivity of the system to variations in the intrinsic cost of plasticity, and iii) the existence of additional factors affecting the evolutionary trajectories. The goal of this analysis is therefore to provide some new insights with the potential to support, refine, and expand previous theoretical findings and perspectives, showing the usefulness of the present ABM as a dynamic tool for generating hypotheses that could also apply to case studies sharing structural similarities with the one illustrated here.

## 2. Methods

### 2.1 Agent-based model

The agent-based model, henceforth referred to as *PhePlastiComp*, was implemented using the modelling environment and programming language NetLogo (Wilensky, 1999) and is freely downloadable at the following URL: https://doi.org/10.5281/zenodo.20452270. The version of NetLogo used in this study is version 6.2.0, that is also available free of charge at the following URL: https://www.netlogo.org/downloads/archive/6.2.0/. However, the model can also be opened using later versions of the software. The user interface of *PhePlastiComp* during a dynamic simulation is shown in Figure 1.

**Figure 1.**
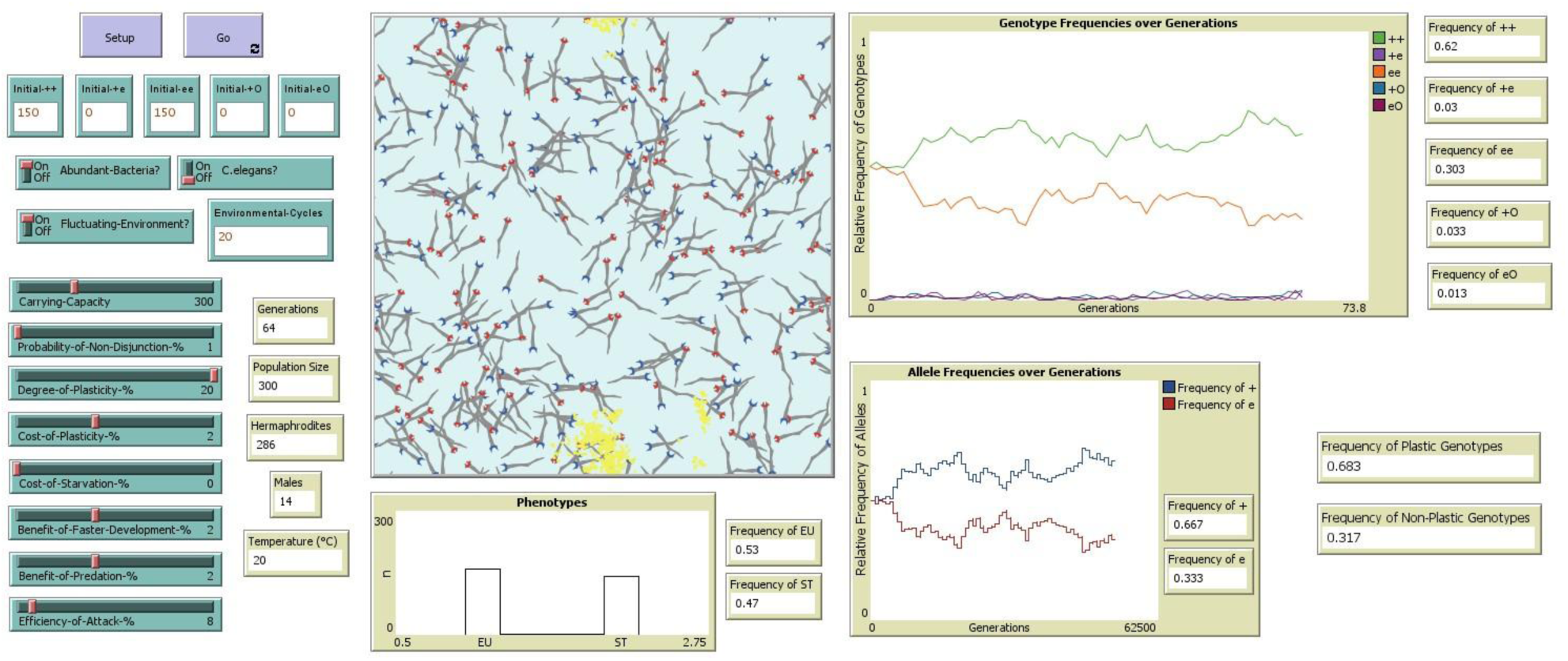
Interface of *PhePlastiComp* during a run. The interface shows the simulation world, that includes a digital culture plate populated by *P. pacificus* nematodes with plastic and non-plastic genotypes and their potential food sources (i.e., *Escherichia coli* bacteria and *Caenorhabditis elegans* larvae), a set of interactive items representing variables and parameters, and several monitors and plots, which are mainly used to keep track of the evolution of the system in terms of allele/genotype/phenotype absolute or relative frequencies.

The model description follows the ODD (Overview, Design concepts, Details) protocol for describing individual- and agent-based models (Grimm et al., 2006; Grimm et al., 2010; Grimm et al., 2020; Grimm et al., 2025), and is included as an appendix to this article (Supplementary File 1). I also provide a further appendix (Supplementary File 2) whose purpose is to introduce all aspects concerning the genetics, development, morphology and ecology of *P. pacificus* that are relevant to providing a more complete context for the implementation of the model. Since all information about *PhePlastiComp* is extensively reported in the above-mentioned files, this section was designed as a concise summary of the latter documents. More specifically, Subsection 2.2 is a very short introduction to *P. pacificus*, Subsection 2.3 is a brief version of the Overview contained within the ODD protocol, as it includes general information on the purpose, agents, and processes of the ABM, and Subsection 2.4 is a concise rationale aimed at providing the biological reasons behind the main modelling choices and parameter values used in the simulated scenarios, also including details on the model validation strategy.

### 2.2 Model organism

*Pristionchus pacificus* is an established model organism in evolutionary biology, mainly used to investigate phenotypic plasticity (Gilarte et al., 2015; Dardiry et al., 2023). This species has a 4-day lifespan at 20°C (Sommer R., 2006) and four different juvenile stages before reaching the adulthood (Hong & Sommer, 2006; Schroeder, 2021). It is identified by an androdioecious mating system, as populations are composed of self-fertilising hermaphrodites (XX) and a small percentage of males (XO) (Pires-daSilva & Sommer, 2004; Morgan et al., 2017).

A key biological feature of *P. pacificus* is its mouth dimorphism, as individual nematodes can develop either an eurystomatous (EU) or a stenostomatous (ST) mouth form irreversibly. Morphological divergences between the two forms lead to alternative dietary behaviours, with EU worms being facultative predators and ST worms exclusively microbial feeders (Sommer R., 2006; Serobyan et al., 2013; Sommer et al., 2017; Sieriebriennikov et al., 2018). Interestingly, both mouth phenotypes are adaptive under specific conditions, as ST nematodes develop faster, particularly in contexts with abundant microbial food, whereas EU nematodes can exploit alternative food sources (e.g., nematodes of other species) when bacteria are scarce (Serobyan et al., 2014; Susoy & Sommer, 2016).

The underlying developmental decision is regulated both stochastically and conditionally (Ragsdale et al., 2013; Susoy & Sommer, 2016; Sommer et al., 2017), so phenotype ratios of a *P. pacificus* population are determined by both chance and a variety of environmental factors including starvation, temperature, and crowding (Bento et al., 2010; Lenuzzi et al., 2021). Most importantly, it is well-acknowledged that mouth-form ratios can be altered by genetic changes, as some dominant, loss-of-function mutations affecting the *eud-1* gene were shown to induce a constitutive expression of the ST phenotype in the lab (Ragsdale et al., 2013; Susoy & Sommer, 2016; Sommer et al., 2017; Sieriebriennikov et al., 2018).

This characterisation makes *P. pacificus* an ideal candidate to explore the evolution of multiple phenotypic strategies, i.e., conditional, stochastic, and constitutive production, within a single framework.

### 2.3 Model overview

The purpose of the model is to simulate the competition between plastic and non-plastic genotypes (depending on the state of the developmental switch, X-linked gene *eud-1* (Ragsdale et al., 2013)) of the dimorphic nematode worm *Pristionchus pacificus* in an in vitro environment with periodic dietary fluctuations. The model allows for the investigation of the effects of several variables – including an intrinsic cost of plasticity, the timescale of environmental fluctuations in terms of non-overlapping generations, and the individual degree of plasticity – on the evolutionary trajectories of populations of *P. pacificus*. The main objective of simulations is to generate data on the frequency and time to fixation of the two competing genotypes and their related alleles. These alleles, hereafter named *+* (wild-type allele) and *e* (mutant allele) are variants of *eud-1*.

The individual entities/agents included in the model are nematodes belonging to two different species, i.e., *P. pacificus* and *Caenorhabditis elegans*, *Escherichia coli* bacteria, and NetLogo patches (grid cells). An additional, high-level agent, the observer, is used to give instructions to all other classes of agents. *P. pacificus* is the main type of agent and is used to explore the competition between plastic and non-plastic genotypes/strains/strategies (see Subsection 2.2 and Supplementary File 2 for more information on this model organism). *C. elegans* is a potential prey of *P. pacificus*, whereas *E. coli* is the main food source of the latter (Serobyan et al., 2014). The set of all patches constitutes the 2D in vitro environment where all these organisms are located, representing a Petri dish or any well-mixed natural environment.

The process executed by the model simulates the entire life cycle of *P. pacificus* nematodes across multiple, non-overlapping generations. Each generation covers 820 discrete time steps (NetLogo ticks), as 10 time steps are used to represent 1 h in the real world and 82 h is approximately the actual lifespan of *P. pacificus* worms (Hong & Sommer, 2006)). The process scheduling is summarised in Figure 2, which also links the functions performed by *P. pacificus* nematodes to the specific time interval when they take place within each generation (i.e., egg and juvenile stages + adult stage).

**Figure 2.**
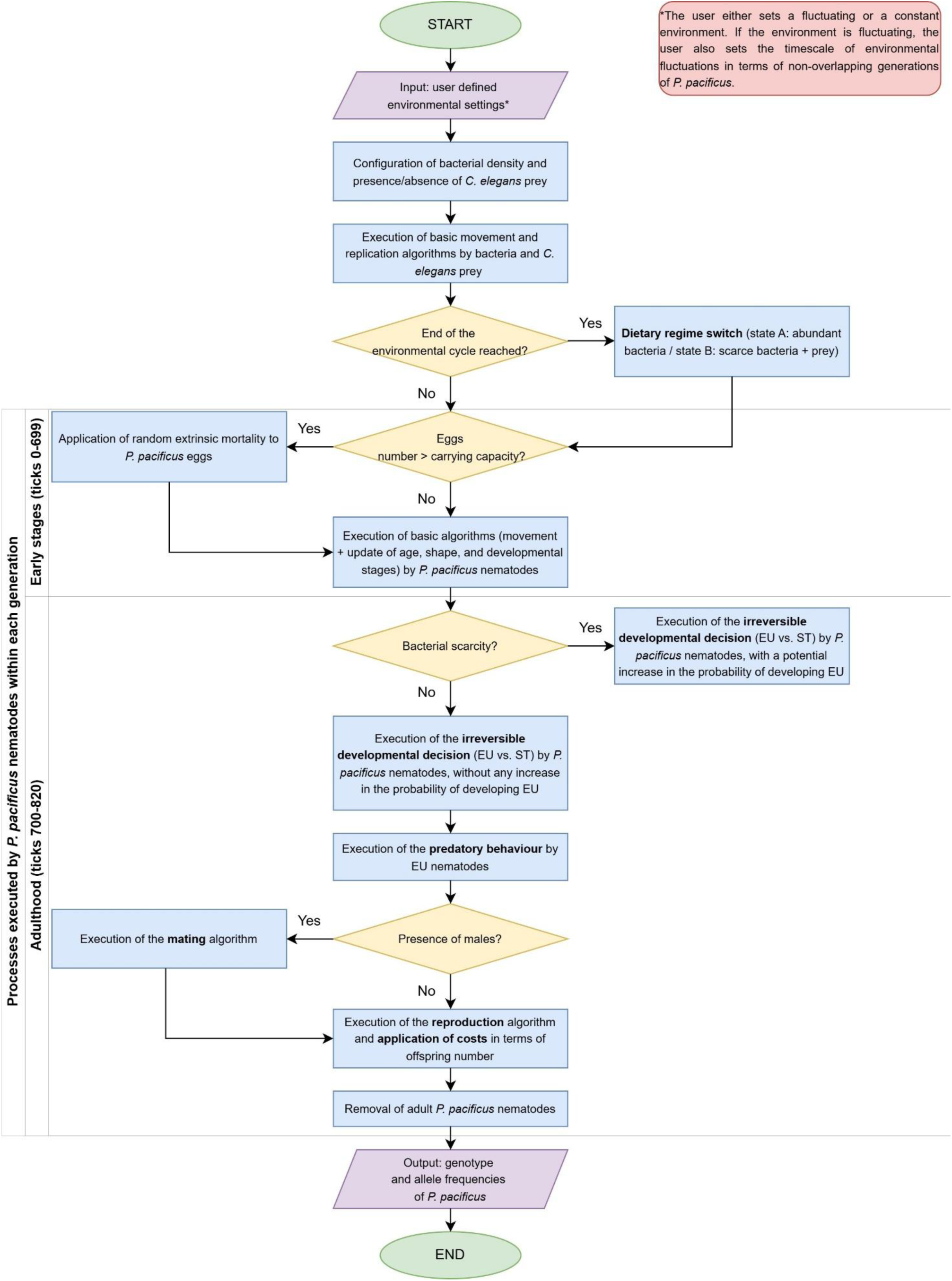
Flowchart of the algorithm. This representation of the process scheduling summarises the steps executed by the model at each discrete time step. It also reports when each function or submodel is executed within *P. pacificus* life cycle, i.e., during the early stages (egg + larval stages) or during the adult stage, as these parts of the program are only activated if worms have reached that specific age/developmental progression interval. The main submodels and critical blocks of the algorithm are highlighted in bold and briefly described in this subsection.

The next paragraphs are dedicated to explaining the key submodels and transitions, strictly following the execution flow of the algorithm. Table 1 summarises the parameters introduced into the system and mentioned in this presentation, also providing a short description of each term, range of values used in my simulations, and references used to parameterise and calibrate the ABM. Additional information is provided in the text below, in Subsection 2.4, and in the ODD protocol (Supplementary File 1).

**Table 1.**
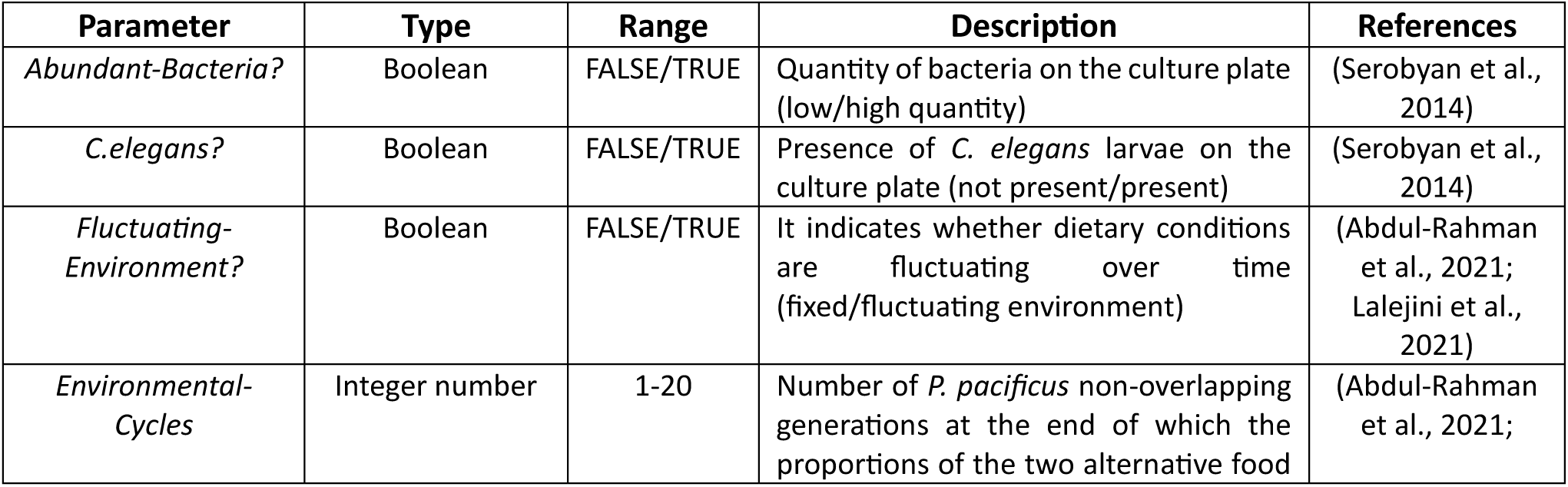
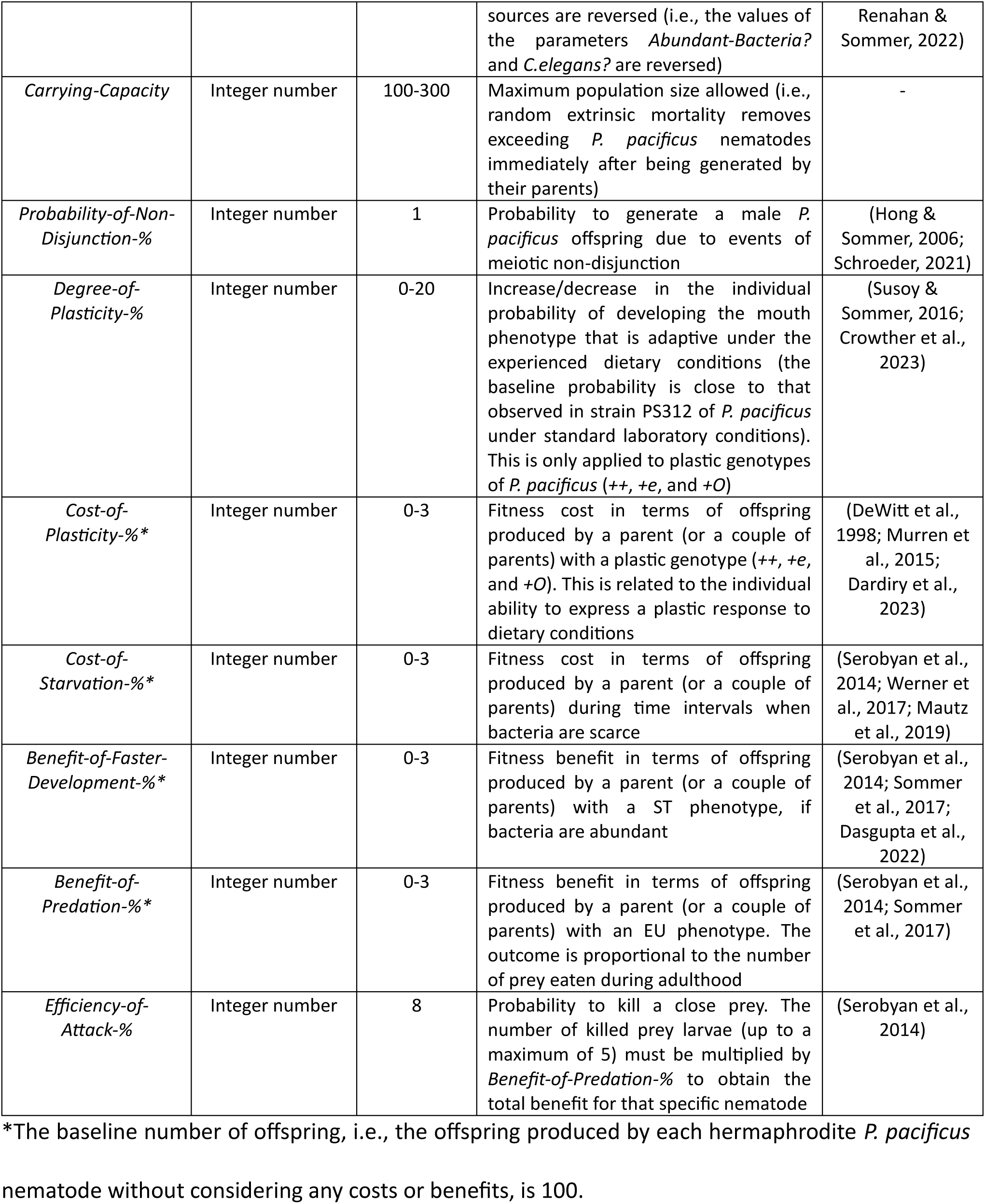
Summary of parameters. Parameters implemented in *PhePlastiComp* and used during simulations. This list includes the type of parameter, the range of values used in the simulated scenarios, a short description, and references supporting the implementation of each parameter and the considered values (if relevant).

Besides initial numbers of *P. pacificus* nematodes for each genotype, i.e., *++*, *+e*, *+O* (plastic), *ee*, and *eO* (non-plastic), the input data are used to inform the model about temporal fluctuations in dietary regimes. Indeed, the environment can be either fixed or fluctuating over time at a specific timescale (expressed in terms of *P. pacificus* generations), with periodic switches of the prevalent nutrient source, i.e., bacteria or *C. elegans* larvae.

The basic movement and replication instructions assigned to food sources are not central, as these secondary agents are used as triggering factors or prey. Conversely, their level of abundance or presence/absence within the environment is a critical component of the system.

The *dietary regime switch* algorithm regulates the transitions between dietary regimes within the virtual environment. If the parameter *Fluctuating-Environment?* = TRUE, the current time step is not 0, and the current time step is a multiple of the product between the duration of a generation of *P. pacificus* (i.e., 820 ticks = 82 hours) and the value of *Environmental-Cycles*, the observer reverses the binary value of *Abundant-Bacteria?* and that of *C.elegans?*. Consequently, the prevalent food source is periodically switched between bacteria and *C. elegans* larvae.

A fixed *Carrying-Capacity* is used to keep population size below a certain value by applying a random extrinsic mortality to the offspring of all *P. pacificus* nematodes immediately after egg laying.

The surviving *P. pacificus* nematodes randomly move, grow, and go through several developmental stages synchronously, i.e., egg, J1, J2, J3, J4, adulthood (J = Juvenile). The most critical transition in their life cycle occurs 700 time steps/ticks (= 70 hours) after oviposition, i.e., at the adult age threshold, as this is when the *irreversible developmental decision* takes place. Depending on their genotype and the value generated by a function that calculates the probability to develop the EU mouth-form, *P. pacificus* nematodes decide to set either the EU (predatory) or the ST (bacterivorous) mouth phenotype. Each genotype has a baseline probability to develop a particular phenotype under standard laboratory conditions, corresponding to a dietary regime with abundant bacteria and no *C. elegans* prey larvae in my runs. The key parameter named *Degree-of-Plasticity-%*, that is only applied to plastic genotypes, is the additional probability to develop the EU morph when bacteria are scarce. Consequently, this parameter enables the regulation of the magnitude of plastic responses to a challenging environment. Of course, nematodes with non-plastic genotypes are not affected by this parameter, as they always develop the ST phenotype. A summary of baseline, genotype-specific mouth-form ratios/probabilities is provided in Table 2, that also shows how and in which contexts *Degree-of-Plasticity-%* is applied.

**Table 2.**
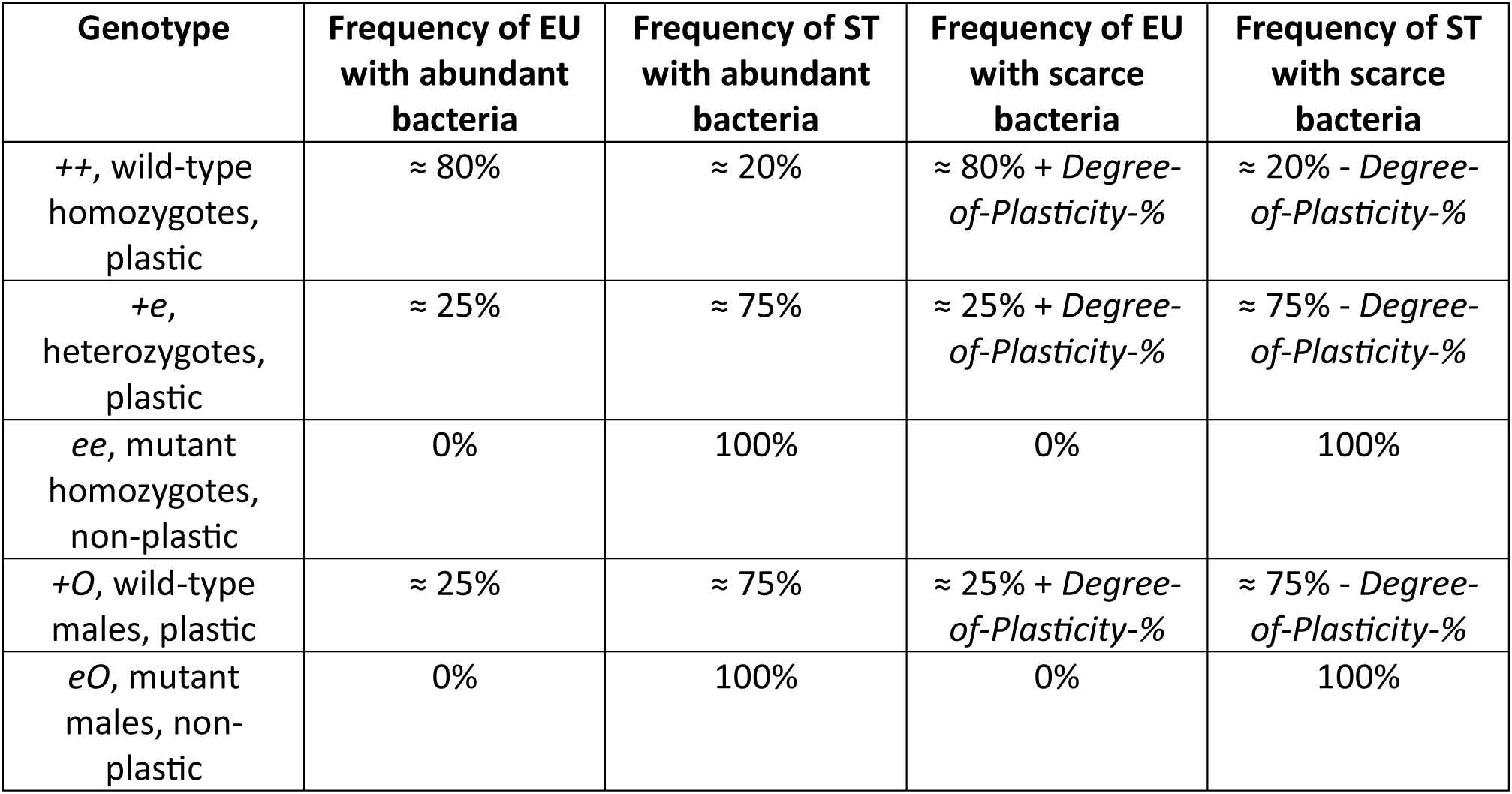
Mouth-form ratios in relation to genotype and variation in food sources.

At this stage of *P. pacificus* life cycle, adult nematodes expressing the EU morph can perform a *predatory behaviour* by eating *C. elegans* larvae (if they are present on the lattice). If an EU worm detects at least one prey within range of a patch, there is a probability corresponding to the value of *Efficiency-of-Attack-%* that this worm kills one of *C. elegans* prey larvae within range of a patch. The number of prey eaten by each EU worm is later used for the calculation of benefits in terms of offspring.

If males are present on the grid, they search for an available hermaphrodite partner (*mating* algorithm): if there are any hermaphrodite nematodes without a partner, each male sets one of those hermaphrodites as a mating partner.

At the end of each generation, the *reproduction* submodel is executed., Offspring are generated by each hermaphrodite worm by following Mendelian segregation ratios (monohybrid selfing or occasional cross, where the locus of interest is the X-linked gene *eud-1*). Considering a baseline offspring number = 100, the total number of offspring (*n_o_*) resulting from self-fertilisation of each digital *P. pacificus* parent is given by Equation (1):

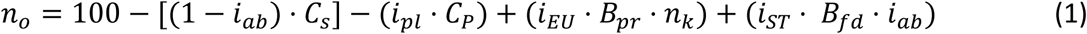

where *C_s_* is the cost of bacterial starvation, *C_P_* is the intrinsic cost of plasticity, *B_pr_* is the benefit in fecundity due to predation, *n_k_* is the number of *C. elegans* larvae killed by the parent, *B_fd_* is the benefit in fecundity due to faster development, and *i_x_* ∈ {0, 1} are Boolean variables indicating environmental or phenotypic states of the parent, i.e., abundance of microbial food (*i_ab_*), plastic genotype (*i_pl_*), EU phenotype (*i_EU_*), and ST phenotype (*i_ST_*).

Benefits in terms of offspring number are applied by each hermaphrodite parent to only one of the resulting filial genotypes, with the same probability of allocation for each genotype. More specifically, *Benefit-of-Faster-Development-%* is only applied to the offspring of ST hermaphrodites (in the event of a crossing, both parents must be ST) if the environment they experience during their life cycle is characterised by microbial abundance, whereas *Benefit-of-Predation-%* is only applied to the offspring of EU hermaphrodites and must be multiplied by the number of prey larvae eaten by parents (both of them, in the event of a crossing) during their life cycle. The value assigned to the parameter *Probability-of-Non-Disjunction-%* determines the probability to produce a male (XO) for each reproductive event.

After reproduction, the *application of costs* algorithm is executed. All costs translate into a removal of a specific number of offspring for each hermaphrodite parent and are randomly applied across filial genotypes. More specifically, *Cost-of-Starvation-%* is due to bacterial scarcity during the life cycle of parents and is applied two times to offspring resulting from cross-fertilisation, whereas *Cost-of-Plasticity-%* is applied to offspring produced by a hermaphrodite with a plastic genotype (this cost is also applied two times if there are two parents and they both have a plastic genotype).

After the costs have been applied to the offspring, all parents immediately die, the new genotype and allele frequencies are reported, and all processes are reiterated for the new generation of nematodes.

### 2.4 Model Parameterisation, biological grounding, and validation

I specifically designed *PhePlastiComp* as a hybrid theoretical-experimental simulation platform. In this regard, the choice of *P. pacificus* as a model organism and the unprecedented focus on full-ST non-plastic strains generated by virtually any hypothetical mutation only affecting mouth-form ratio was crucial. In addition to providing a context for producing preliminary data on still unexplored competition dynamics – indeed, despite the recent discovery of a full-ST natural isolate (see interview with Ralf J. Sommer (2025)), previous assessments have necessarily used wild, full-EU non-plastic strains (Dardiry et al., 2023; Kalirad & Sommer, 2024) – *P. pacificus* could hopefully be used as a proxy for specific mixed-strategy dimorphic systems (see Subsection 3.6 on model applicability).

Consequently, strategies of model parameterisation and validation were oriented at ensuring that the computational system remained mostly grounded in the currently available information on the modelled species while enabling the exploration of the parameter space using sufficiently broad, yet ecologically realistic ranges of key parameter values.

In line with these principles, the ABM relies on both fixed parameter values derived from the literature and parameters subjected to systematic sweeps. While extensive biological information underlying all parameters and processes included in *PhePlastiComp* is detailed in the ODD protocol (Supplementary File 1), I use the following points to illustrate the rationale behind the main parameters and ranges chosen for my exploratory analysis:

- **Costs and benefits**: Costs and benefits in fitness/fecundity ranging from 0% up to 3% – depending on the considered scenario, see Subsection 2.5 – were chosen based on a principle of ecological plausibility, as these represent conservative parameter estimates, avoiding unrealistic regimes. As for the logic underlying the application of costs of plasticity only to *++*, *+e*, and *+O* genotypes, this procedure is consistent with observations conducted with mutagenised *P. pacificus* nematodes (Ragsdale et al., 2013) showing that a single mutation cannot completely defuse the molecular machinery responsible for the developmental decision, so any inherent plasticity costs would likely persist.
- **Carrying capacity**: Carrying capacities ranging from 100 to 300 nematodes – whose effects on the system have been assessed in a sensitivity analysis focusing on an exemplary scenario, see Subsection 3.3 – were selected to exclude any condition of overcrowding, which is considered an additional factor also affecting mouth-form ratios in this species (Bento et al., 2010). However, carrying capacity was generally kept fixed at 300 to achieve a balance between computational feasibility and minimisation of potential noise due to genetic drift. Moreover, limited population sizes may match better a physically bounded environment where variation is most likely temporal rather than spatial, in line with model’s design.
- **Degree of plasticity**: Degrees of plasticity ranging from 0% to 20% guarantee systematic exploration of the effects of variation in this key parameter, including values approximating 0% (close to exclusive stochastic regulation of plasticity), moderate (10%) and relatively high (20%) levels of plasticity. The upper limit of 20% was chosen to match the maximum real value potentially evolvable by PS312 *P. pacificus* strain, as adding this percentage to the strain-specific baseline proportion of EU = 80% (Susoy & Sommer, 2016) would induce the production of 100% EU worms under starved conditions. Intraspecific variation in the degree of plasticity is common and has been documented in many taxa, from chordates to plants (Walls, 1997; Nussey et al., 2005; Lind & Johansson, 2009; Bhat et al., 2015; Grewell et al., 2016). Since this degree is plausibly controlled by gene regulatory networks (e.g., the pathway including *eud-1* as a key node), natural selection can affect it (Susoy & Sommer, 2016), which legitimates a theoretical exploration aimed at testing the performance of a set of genotypes that differ in their phenotypic responsiveness to environmental cues.
- **Environmental cycles**: Timescales of environmental fluctuations ranging from 1 to 20 *P. pacificus* generations were selected because: i) between generations changes are a reasonable assumption for an organism whose life cycle includes an irreversible developmental decision (Kassen, 2002), ii) roughly periodic changes may reflect the real trends of a wide range of biological systems, which are often characterised by periodic environmental oscillations at the most disparate scales (Abdul-Rahman et al., 2021), and iii) in addition to covering fast, intermediate, and slow timescales, *Environmental-Cycles* = 1, 5, 10, 15, and 20 could match data obtained from real ecosystems where *P. pacificus* can be found. For instance, in carcasses of beetles co-infested by other nematodes, mouth-form ratios can undergo major changes within a few weeks (Renahan & Sommer, 2022), which is consistent with environmental variation across several generations of worms. Considering the heterogeneity of *P. pacificus* habitats, which include soil, scarab bodies and corpses, and decomposing vegetal matter (Susoy & Sommer, 2016; Félix et al., 2018), it is also plausible that, in some cases, a specific nutrient source may be more prevalent than another (on average) for a time interval spanning several generations.

After parameterisation, the model underwent a hierarchical validation process aimed at assessing its reliability both at the agent- and system-level. Micro-validation (agent level) was conducted by comparing individual rules and parameters to data drawn from the relevant literature on this organism (Supplementary Files 1 and 2). Macro-validation (system level) followed a Pattern-Oriented Modelling (POM) approach (Grimm, 2005), integrating preliminary scenarios explicitly designed to test the ability of the model to produce basic expected outcomes (Supplementary File 3), and main scenarios conceived to assess the matching between simulation outcomes and established dynamics – either reported in experimental evolution studies or derived from general theory – while searching for potentially new trends (see Subsections 3.1 and 3.2).

### 2.5 Simulated scenarios

Since agent-based models rely on large numbers of computational individuals whose interactions are often regulated by probabilistic instructions, the final outcomes resulting from a single run are expected to differ from those generated by another run sharing the same settings (Hunter & Kelleher, 2020). For this reason, all results produced by each simulated scenario are extracted from multiple runs, more specifically, 100 or 300 runs for Symmetric Analyses (SAs) and 1000 runs for Invasibility Analyses (IAs). Symmetric Analyses start with an equal number of *P. pacificus* nematodes belonging to two alternative strains (i.e., plastic vs non-plastic), whereas Invasibility Analyses start with a single mutant or immigrant individual belonging to a strain different from that of the population to be invaded.

Preliminary scenarios are used to validate qualitatively the model at a system level (macro-validation), based on the match between the outcomes of runs and some expected patterns that can be inferred from the relevant literature (see Table 3A). The outcomes of these preliminary scenarios are presented in Supplementary File 3. Additional validation at a system level is also provided by outcomes generated by the main simulated scenarios, as they also consistently reproduced patterns in line with empirical evidence and pre-existing theoretical models regarding the evolution of plasticity (see Subsections 3.1 and 3.2).

**Table 3A.**
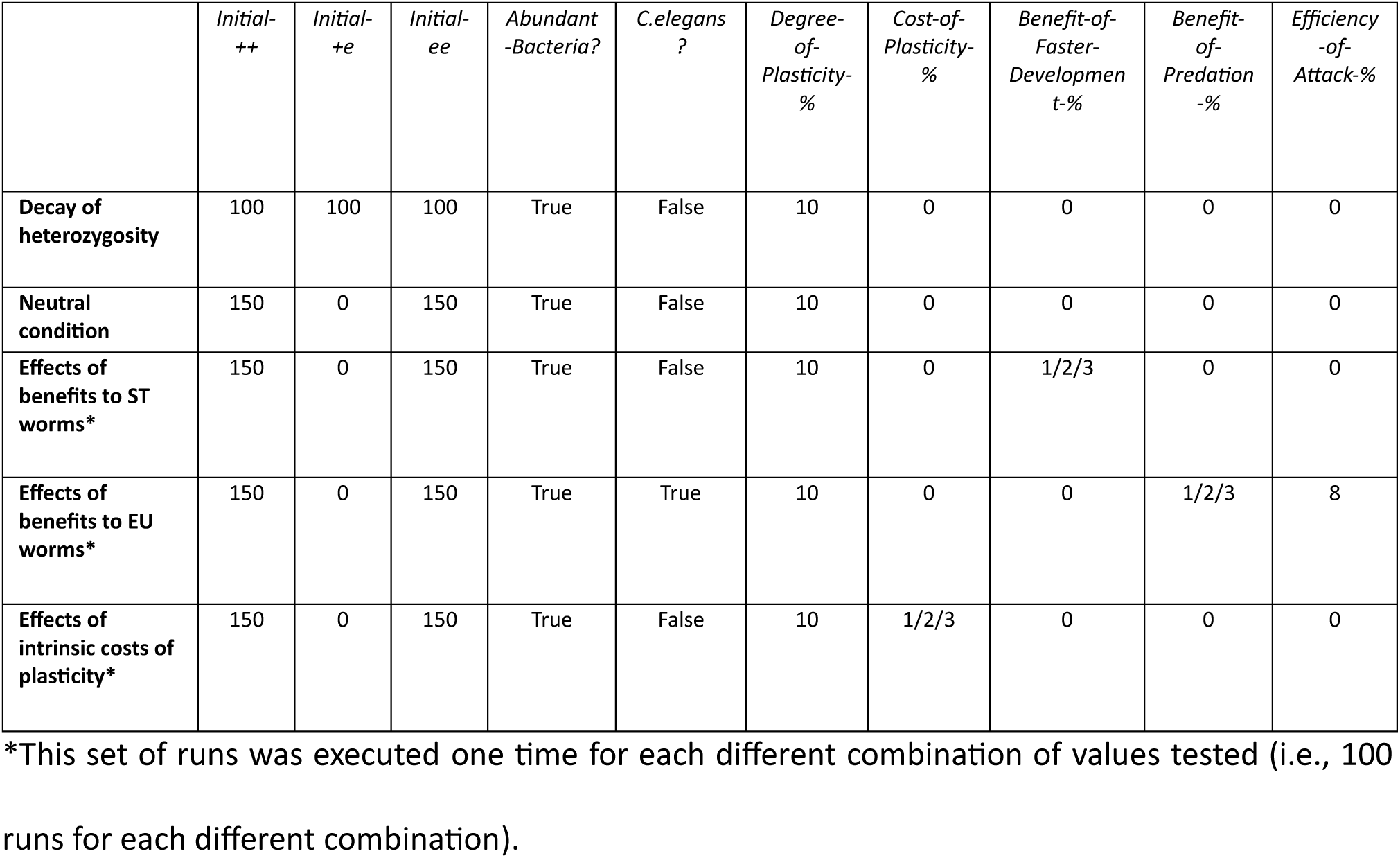
Validation scenarios. List of all parameter values used for each scenario aimed at validating the global performance of the simulated system. In all these scenarios, *Fluctuating-Environment* = False, *Environmental-Cycles* = N/A, *Carrying-Capacity* = 300, *Probability-of-Non-Disjunction-%* = 0, and *Cost-of-Starvation-%* = 0.

|  | *Initial-*<br>*++* | *Initial-*<br>*+e* | *Initial-*<br>*-ee* | *Abundant*<br>*-Bacteria?* | *C.elegans*<br>*?* | *Degree-*<br>*of-*<br>*Plasticity-*<br>*%* | *Cost-of-*<br>*Plasticity-*<br>*%* | *Benefit-of-*<br>*Faster-*<br>*Developmen*<br>*t-%* | *Benefit-*<br>*of-*<br>*Predation*<br>*-%* | *Efficiency*<br>*-of-*<br>*Attack-%* |
| --- | --- | --- | --- | --- | --- | --- | --- | --- | --- | --- |
| Decay of heterozygosity | 100 | 100 | 100 | True | False | 10 | 0 | 0 | 0 | 0 |
| Neutral condition | 150 | 0 | 150 | True | False | 10 | 0 | 0 | 0 | 0 |
| Effects of benefits to ST worms* | 150 | 0 | 150 | True | False | 10 | 0 | 1/2/3 | 0 | 0 |
| Effects of benefits to EU worms* | 150 | 0 | 150 | True | True | 10 | 0 | 0 | 1/2/3 | 8 |
| Effects of intrinsic costs of plasticity* | 150 | 0 | 150 | True | False | 10 | 1/2/3 | 0 | 0 | 0 |

\*This set of runs was executed one time for each different combination of values tested (i.e., 100 runs for each different combination).

The main scenarios explored in this study were used to investigate the combined effects of several parameters, including the intrinsic cost of plasticity, the benefits associated to each phenotype, the degree of plasticity, and the timescale of environmental fluctuations, on the evolutionary interplay between plastic and non-plastic strains (see Table 3B). This interplay was monitored using relative frequencies and times to fixation (in terms of non-overlapping generations) of the alleles *+* and *e*. Note that all simulations started with only hermaphrodite nematodes, that is, *Initial-+O* and *Initial-eO* are always set to 0 in time step 0 (i.e., NetLogo tick = 0). In each fluctuating scenario, runs were split evenly: half started with abundant bacteria and no prey, the other with scarce bacteria and prey.

**Table 3B.** Scenarios explored. List of all parameter values used for each hypothetical scenario explored in this study. In all these scenarios, *Carrying-Capacity* = 300, *Probability-of-Non-Disjunction-%* = 1, *Cost-of-Starvation-%* = 3, and *Efficiency-of-Attack-%* = 8.

|  | *Initial*<br>*-++* | *Initial*<br>*-+e* | *Initial*<br>*-ee* | *Abunda*<br>*nt-*<br>*Bacteria*<br>*?* | *C.elegan*<br>*s?* | *Fluctuatin*<br>*g-*<br>*Environme*<br>*nt?* | *Environmen*<br>*tal-Cycles* | *Degree-*<br>*of-*<br>*Plasticit*<br>*y-%* | *Cost-of-*<br>*Plasticit*<br>*y-%* | *Benefit-of-*<br>*Faster-*<br>*Developm*<br>*ent-%* | *Benefit-*<br>*of-*<br>*Predatio*<br>*n-%* |
| --- | --- | --- | --- | --- | --- | --- | --- | --- | --- | --- | --- |
| SA:<br>Impact<br>of<br>environmenta<br>l<br>change<br>and<br>degree<br>of plasticity* | 150 | 0 | 150 | True/Fal<br>se | False/Tr<br>ue | True | 1/5/10/15/<br>20 | 0/10/2<br>0 | 0/1 | 2 | 2 |
| SA:<br>Tolerance<br>of<br>plastic<br>strains<br>towards<br>the cost<br>of plasticity* | 150 | 0 | 150 | True/Fal<br>se | False/Tr<br>ue | True | 20 | 20 | 0/1/2/3 | 2 | 2 |
| IA: Impact of cost of plasticity on the fixation of non-plastic strains* | 299 | 1 | 0 | True | False | False | N/A | 10 | 0/1/2/3 | 2 | 2 |
| IA: Impact of differences between the two benefits* | 0/1 | 0/1 | 299 | True/False | False/True | True | 10 | 0/20 | 0/1 | 1 | 1/2/3 |

\*This set of runs was executed one time for each different combination of values tested. Specifically, 100 runs were executed for each different combination in SA scenarios, except for combinations of the SA “Impact of environmental change and degree of plasticity” with 1% cost of plasticity, which were run 300 times. 1000 runs were executed for each different combination of values in IA scenarios.

## 3. Results and Discussion

### 3.1 Higher plasticity and moderate-to-long environmental cycles may be weak promoters of plastic strategies

Symmetric Analyses were carried out using not very likely starting conditions, that is, equal numbers of individuals for both the alternative strains involved (Kalirad & Sommer, 2024) (*++* and *ee*). However, this simplification makes it possible to identify several interesting population dynamics and explore the global impact of the main variables that were introduced into the system. From now on, the symbols *ƒ_fix_(+)* and *ƒ_fix_(e)* are used to denote the relative frequency of fixation of the two competing alleles, while the symbol *t̅_fix_(e)* is used to denote the mean time until fixation of the mutant allele.

The aim of the SA named “Impact of environmental change and degree of plasticity” is to explore the advantages of plastic strains with a different level of plasticity across several timescales of environmental fluctuation. To do this, I report the relative frequencies of the alleles *+* and *e* extracted from runs executed with the following initial settings: *Initial-++* = 150, *Initial-ee* = 150, *Carrying-Capacity* = 300, alternate presence of abundant bacteria and *C. elegans* prey larvae (= fluctuating environment), *Environmental-Cycles* = 1, 5, 10, 15, or 20, *Probability-of-Non-Disjunction-%* = 1, *Degree-of-Plasticity-%* = 0, 10, or 20, *Cost-of-Plasticity-%* = 0 or 1, *Cost-of-Starvation-%* = 3, equal benefits in fecundity for both mouth phenotypes = 2 (i.e., *Benefit-of-Faster-Development-%* = *Benefit-of-Predation-%* = 2), and *Efficiency-of-Attack-%* = 8. The different timescales of environmental fluctuations (*Environmental-Cycles*) are the non-overlapping generations of *P. pacificus* it takes before the prevalent nutrient sources available within the NetLogo lattice, i.e., bacteria or *C. elegans* larvae, are reversed.

I first report the outcomes obtained with no costs associated with the production and maintenance of plastic responses (i.e., *Cost-of-Plasticity-%* = 0) across three alternative versions of this scenario including different degrees of plasticity: i) 0% (stochastic regulation without sensitivity to the environment), ii) 10%, and iii) 20%. This SA was carried out using 100 runs for each different combination of environmental periods and levels of plasticity. The outcomes suggest a positive effect of increasing levels of plasticity on *ƒ_fix_(+)* and a plausible role of the duration of environmental cycles in influencing competition dynamics (Figure 3). Specifically, moderate-to-long periods of environmental stability – ranging from 10 to 20 generations – might increase *ƒ_fix_(+)* in all configurations, although this advantage seems to decrease when nematodes are subjected to the longest timescale considered here (20, see Figure 3A and 3B) and is not clearly observed when using *Degree-of-Plasticity-%* = 20 (Figure 3C).

**Figure 3.**
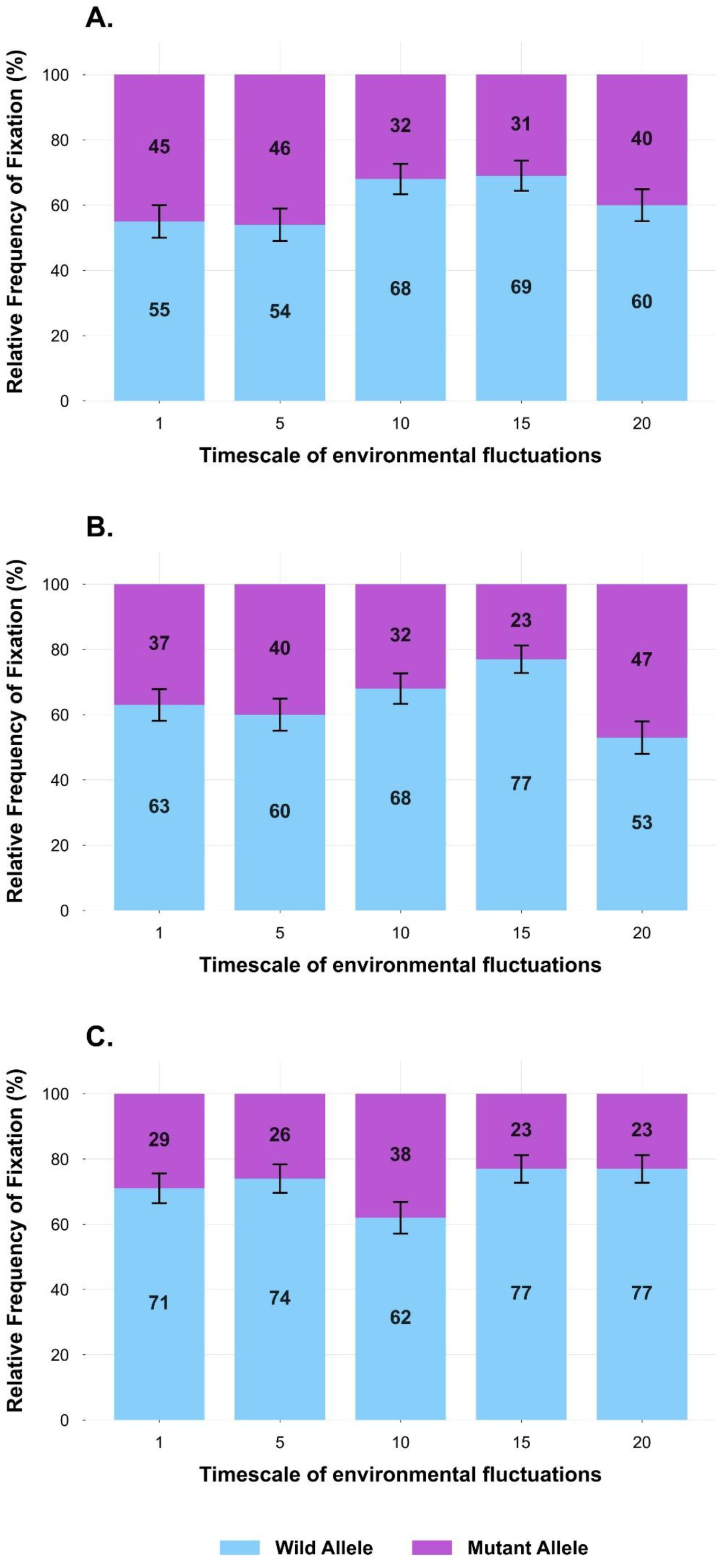
Effects of different timescales of environmental fluctuation on the relative frequencies of fixation of the two competing alleles with no intrinsic cost of plasticity. Plots were constructed using data generated by the SA named “Impact of environmental change and degree of plasticity”. In this version of the scenario, inherent costs of plasticity are absent. Stacked bars represent frequencies of fixation of the two competing alleles (percentages) across five alternative timescales of environmental fluctuations, i.e., 1, 5, 10, 15, and 20, corresponding to numbers of non-overlapping *P. pacificus* generations. Numbers displayed within segments represent the percentage of fixation of each allele, and error bars indicate the standard error (SE) of binomial proportions. In Panel **A**, the plastic response is only stochastically regulated (i.e., *Degree-of-Plasticity-%* = 0). In Panel **B**, outcomes are generated introducing a moderate degree of plasticity (= 10%). Panel **C** shows outcomes produced with the highest degree of plasticity available in *PhePlastiComp* (= 20%). In all panels, light blue = *+* and purple = *e*. The total number of runs executed for each alternative timescale was 100.

As expected by visual inspection of data, logistic regression analysis revealed a significant positive relationship between the degree of plasticity and fixation rates of the wild-type allele (β = 2.46, *z* = 3.66, *p* < 0.001).

The relationship between timescales of fluctuations and frequency of fixation of the wild-type allele (*+*) was formally analysed using a logistic regression model including both a linear and a quadratic term for environmental-cycle duration, to capture the possible non-monotonic trend observed in simulation data. When using a null degree of plasticity (Figure 3A), the linear term showed an *OR* = 1.12 (95% *CI*: 1.01-1.24, *p* = 0.03), whereas the quadratic term showed an *OR* = 0.996 (95% *CI*: 0.99-1, *p* = 0.08). While the significance of the linear component confirms an overall increase of *ƒ_fix_(+)* in response to increasing duration of cycles, the quadratic component might also suggest a marginal curvilinear trend, consistent with the pattern observed across the five environmental conditions, where *ƒ_fix_(+)* tends to decrease with *Environmental-Cycles* = 20.

A clearer demonstration of the same trend is provided by the version of this scenario including an intermediate value of *Degree-of-Plasticity-%* (= 10), as the linear term showed an *OR* = 1.14 (95% *CI*: 1.02-1.26, *p* = 0.02) and the quadratic term an *OR* = 0.99 (*CI* 95%: 0.99-1, *p* = 0.01). The significance of both terms indicates a clear non-monotonic, concave trend, consistent with the pattern suggested by Figure 3B.

In contrast, while producing the highest number of fixations of *+* in most environmental conditions – especially when using the longest durations of environmental cycles – the third version of this cost-free scenario does not provide additional confirmation of the pattern, as the linear term showed an *OR* = 0.94 (95% *CI*: 0.84-1.06, *p* = 0.32) and the quadratic term an *OR* = 1.003 (95% *CI*: 1-1.01, *p* = 0.19).

Collectively, these outcomes obtained in a cost-free condition suggest two trends: i) a positive relation between the frequency of fixation of the *+* allele and the level of plasticity, and ii) an influence of environmental fluctuations in competition dynamics. Indeed, intermediate-to-long environmental cycles might enhance the probability of fixation of *+* in different conditions, i.e., with plastic genotypes adopting either pure bet-hedging or a combination of stochastic and conditional regulation of plasticity.

To determine whether the same timescale-dependent patterns would emerge when introducing an inherent cost of plasticity, I performed again this SA using the same settings, except for the introduction of a low cost of plasticity (= 1%). These versions of the scenario were generated using 300 runs for each different combination of levels of plasticity and environmental cycles, as the number of fixations of *+* using 100 runs was too low to enable a proper assessment of the system’s behaviour. The observed outcomes suggest an increased impact of stochastic oscillations in the simulation process, as some slight emerging tendencies could only be detected when using the highest value of *Degree-of-Plasticity-%* (= 20) (see Figure 4).

**Figure 4.**
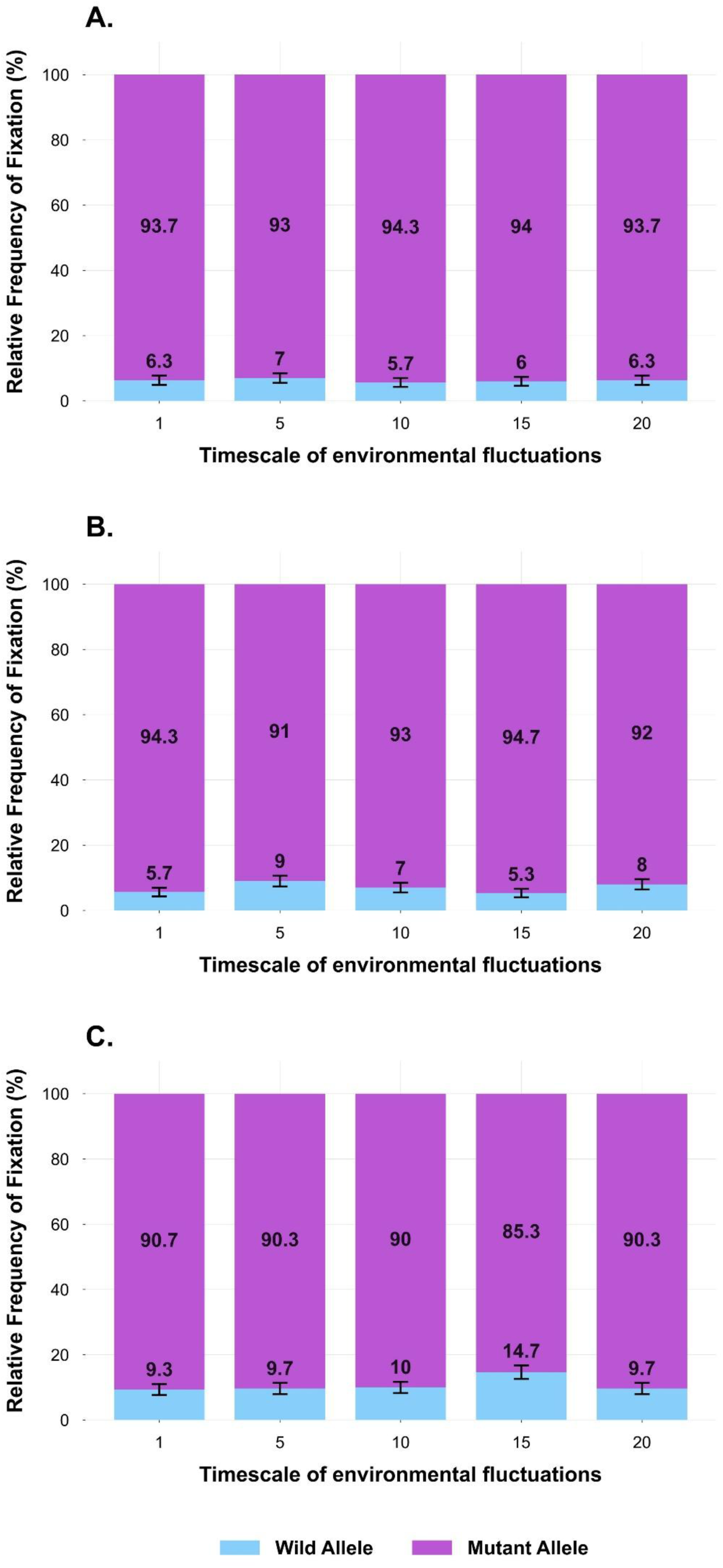
Effects of different timescales of environmental fluctuation on the relative frequencies of fixation of the two competing alleles with a low intrinsic cost of plasticity. Plots were constructed using data generated by the SA named “Impact of environmental change and degree of plasticity”. This version of the scenario includes an inherent, low cost of plasticity (= 1%). Stacked bars represent frequencies of fixation of the two competing alleles (percentages) across five alternative timescales of environmental fluctuations, i.e., 1, 5, 10, 15, and 20, corresponding to numbers of non-overlapping *P. pacificus* generations. Numbers displayed within segments represent the percentage of fixation of each allele, and error bars indicate the standard error (SE) of binomial proportions. In Panel **A**, the plastic response is only stochastically regulated (i.e., *Degree-of-Plasticity-%* = 0). In Panel **B**, outcomes are generated introducing a moderate degree of plasticity (= 10%). Panel **C** shows outcomes produced with the highest degree of plasticity available in *PhePlastiComp* (= 20%). In all panels, light blue = *+* and purple = *e*. The total number of runs executed for each alternative timescale was 300.

The application of quadratic logistic regression to the first two versions of this scenario including a cost of plasticity (i.e., with exclusively stochastic regulation of plasticity and 10% degree of plasticity, respectively) strongly suggests the absence of a trend comparable to that uncovered by simulations performed in a cost-free context. More specifically, the non-significance of both the linear term (*OR* = 0.98, 95% *CI*: 0.87-1.11, *p* = 0.77) and the quadratic term (*OR* = 1, 95% *CI*: 0.99-1.01, *p* = 0.81) when considering stochastic regulation of plasticity alone (Figure 4A), and the non-significance of both terms when adding a level of environmental sensitivity (linear term: *OR* = 1, 95% *CI*: 0.89-1.13, *p* = 0.95; quadratic term: *OR* = 1, 95% *CI*: 0.99-1.005, *p* = 0.98, see Figure 4B) indicate that, in these conditions, the duration of environmental cycles does not affect competition dynamics.

Interestingly, the third version of this costly scenario (i.e., the version using *Degree-of-Plasticity-%* = 20) suggests that the emergence from noise of the trends observed in the cost-free system might only start occurring when the level of conditional regulation of the plastic response is above a specific threshold (in the model, between 10% and 20%). While both the linear term (*OR* = 1.07, 95% *CI*: 0.97-1.18, *p* = 0.19) and the quadratic term (*OR* = 0.997, 95% *CI*: 0.99-1, *p* = 0.27) are still non-significant, the decrease in p-values compared to the previous tests might reinforce the visual insights provided by Figure 4C, indicating that the period of environmental stability could still be a weak modulator of the system, as intermediate-to-long environmental cycles could provide a slight advantage to plastic strategists (an advantage which might decrease with longer cycles). Simultaneously, *ƒ_fix_(+)* could start displaying a general increase – compared to the two previous versions of this SA including a cost of plasticity – due to the higher degree of plasticity assigned to plastic genotypes.

This SA provides both confirmations of expected patterns, which are valuable from a macro-validation perspective, and novel insights deserving further investigation. First, the outcomes obtained under cost-free conditions clearly indicate that plastic genotypes exhibiting higher developmental sensitivity are favoured by selection in a regularly fluctuating environment. Specifically, when plasticity is not costly, while an intrinsic advantage is provided by the constant diversification of phenotypes in both dietary regimes, which confirms the role of stochasticity as a bet-hedging strategy in PS312, this baseline benefit is powered by environmental responsiveness, that further increases the fraction of individuals developing the optimal phenotype under each diet.

This emerging pattern generated by *PhePlastiComp* is fully consistent with the outcomes of experimental evolution studies involving another nematode species combining bet-hedging and plasticity, i.e., *Caenorhabditis remanei* (Sekajova et al., 2025). Indeed, when subjected to a regime consisting of regular between-generations temperature switches, worms evolved increased phenotypic plasticity in body size, implying that more plastic genotypes were favoured under fluctuating environmental conditions. While the sharing of this evolutionary outcome provides relevant empirical support to the model from a macro-validation perspective, it is also important to notice that the above analysed results could also expand the scope of this experimental study, as the positive relationship between degree of plasticity and frequency of fixation of plastic strains seems to be not limited to environments fluctuating at a fast rate (as is the case with the study using *C. remanei*).

Interestingly, the finding that a moderate-to-long duration of alternative dietary regimes may increase the advantage of plastic strains in *P. pacificus* could also be supported by (and provide support to) recent theoretical results obtained with a partially comparable mathematical model based on the Lotka-Volterra equation and focusing on the competition between one plastic generalist and two specialist genotypes (clones) of an asexually reproducing species (Kasada & Yoshida, 2020). In this context, “generalist” and “specialist” are used as synonyms of “plastic” and “non-plastic”, respectively, although the former couple of terms primarily refers to the concept of fitness under different environmental conditions and is not exclusively related to the notion of plasticity (van Tienderen, 1997). The outcomes of the mathematical model suggest that both on moderate and long timescales of environmental fluctuations (in the latter case, only if the cost of plasticity is sufficiently low) the population with plasticity may increase in frequency and, therefore, the generalist genotype may dominate over the specialist one.

As argued by the authors, a possible explanation for this pattern is that the specialist genotype that experienced the maladaptive environment for a sufficiently extended period requires a long time to recover its density during the next environmental cycle. While *PhePlastiComp* also includes a stochastic component in phenotype production, this explanation is also compatible with the overall dynamics reported in this SA: when the type of food available changes at a slower pace, non-plastic sub-populations periodically face a longer “desert crossing” during which they are severely affected by starvation and do not experience any benefits, whereas plastic sub-populations can exploit the benefits of each environment by developing a variable fraction of the optimal morph.

Furthermore, in the ABM, there seems to be an upper temporal limit beyond which these benefits for plastic sub-populations diminish, an additional pattern that, in the analytical model, was only observed when using non-negligible costs of plasticity. Intriguingly, analogous suggestions on the role of intermediate timescales in selecting generalist strategies are also found outside evolutionary biology, e.g., in affinity maturation, the process through which the immune system produces high-affinity antibodies for virtually any antigen (Mishra & Mariuzza, 2018; Sachdeva et al., 2020).

While the closeness between the outcomes presented here and those illustrated in Kasada & Yoshida (2020) further supports the reliability of *PhePlastiComp* through model cross-validation, the introduction of stochasticity and different degrees of plasticity may reveal some unprecedented details. Indeed, SA configurations including 1% cost of plasticity suggest that the influence of environmental fluctuation rates might only be relevant when the magnitude of plastic responses is above a minimum threshold, which would make the duration of environmental periods a weak evolutionary modulator of strategies combining phenotypic plasticity and bet-hedging. However, further research is needed to test this hypothesis and possibly identify a threshold level of plasticity beyond which the fitness benefits for plastic genotypes start emerging from noise, especially using alternative approaches and model organisms.

Assuming a positive relationship between degree and cost of plasticity – which has been hypothesised by theoretical studies and supported by empirical research (van Tienderen, 1997; Lind & Johansson, 2009) – these outcomes could support the speculation that plastic genotypes of organisms sharing the main biological characteristics of *P. pacificus* might be selected if their degree of conditional plasticity is either i) very low (or = 0), approximating performances displayed in the cost-free scenario only involving bet-hedging (Figure 3A), or ii) high enough to “unlock” relevant benefits provided by moderate-to-slow environmental fluctuations and greater matching with the environment, provided that the cost function remains sufficiently shallow.

### 3.2 Increasing costs of plasticity might induce a two-step phase transition by preventing fixation of plastic genotypes and reducing coexistence

The SA named “Tolerance of plastic strains towards the cost of plasticity” is based on the following settings: *Initial-++* = 150, *Initial-ee* = 150, *Carrying-Capacity* = 300, alternate presence of abundant bacteria and *C. elegans* prey larvae (= fluctuating environment), *Environmental-Cycles* = 20, *Probability-of-Non-Disjunction-%* = 1, *Degree-of-Plasticity-%* = 20, *Cost-of-Plasticity-%* = 0, 1, 2, and 3, *Cost-of-Starvation-%* = 3, equal benefits in fecundity for both mouth phenotypes = 2 (i.e., *Benefit-of-Faster-Development-%* = *Benefit-of-Predation-%* = 2), and *Efficiency-of-Attack-%* = 8. The relatively high level of plasticity and timescale of environmental change were selected because it turned out from the previous SA that, in some cases, these settings may increase the probability of fixation of the wild-type allele, so they could possibly mitigate the negative effects provided by the cost of plasticity.

Contrary to this expectation, this scenario informs that even moderate costs of plasticity may be sufficient to prevent the fixation of the *+* allele. Indeed, *ƒ_fix_(+)* dropped sharply from 77% (cost-free condition) to 14% (*Cost-of-Plasticity-%* = 1), while no fixation events occurred when using costs ≥ 2% (see Figure 5A). While this abrupt collapse in the frequency of fixation seems to indicate the existence of a tipping point between 0% and 1% cost, times until fixation of the mutant allele could show a second phase transition between 1% and 2% cost. This is suggested by a relevant reduction in the dispersion of data and by the progression of average times to fixation, as *t̅_fix_(e)* = 469.74, 333.07, 180.09, and 130.97 when using a *Cost-of-Plasticity-%* = 0, 1, 2, and 3, respectively (Figure 5B).

**Figure 5.**
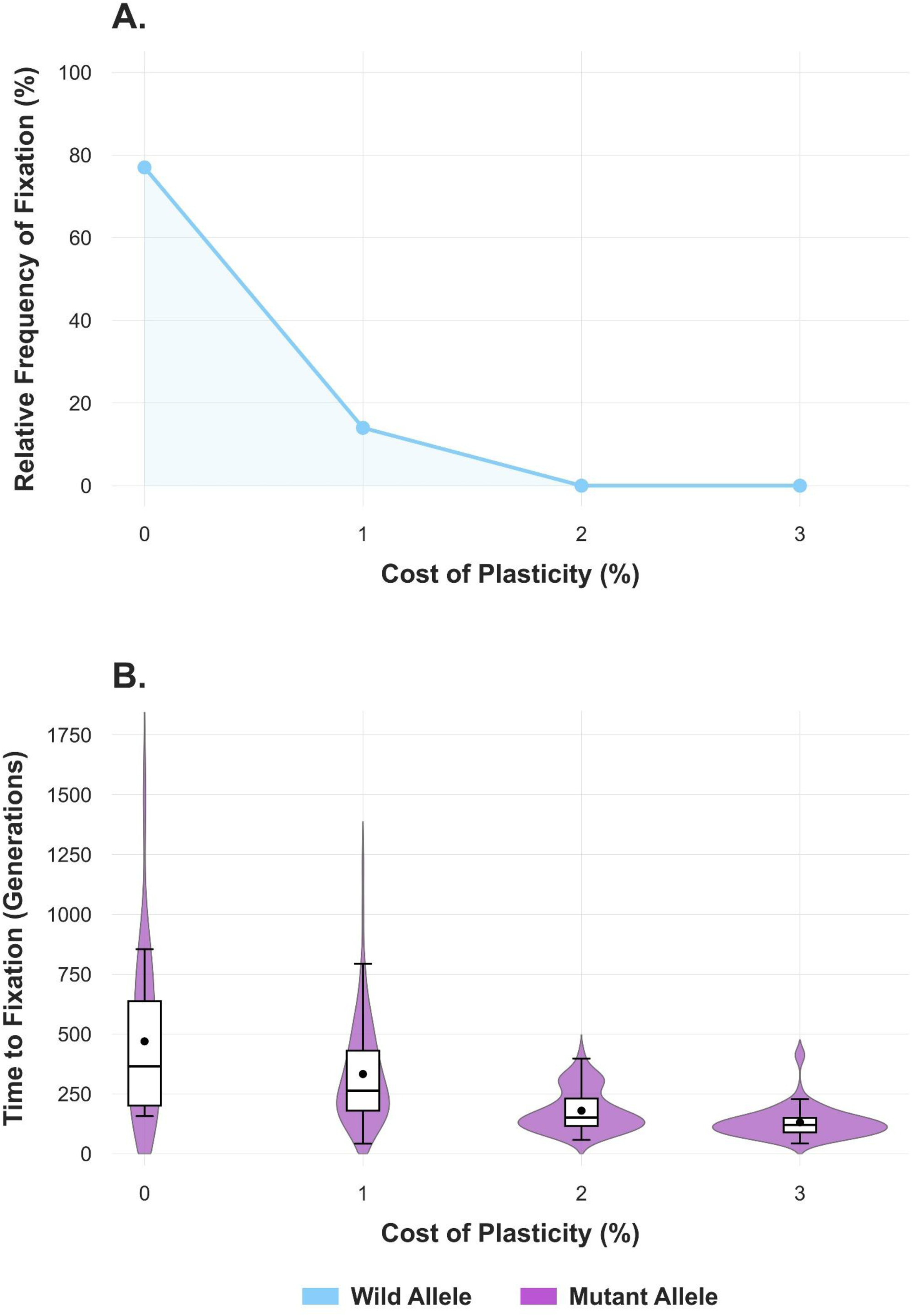
Effects of increasing costs of plasticity on the relative frequency of fixation of the wild-type allele and time until fixation of the mutant allele. These plots were constructed using data generated by the SA “Tolerance of plastic strains towards the cost of plasticity”. In Panel **A**, a line plot shows the effects of increasing costs of plasticity on the relative frequency of fixation of the wild allele. In Panel **B**, violin plots with overlaid boxplots show the distributions of times to fixation of the mutant allele – expressed in terms of non-overlapping *P. pacificus* generations – for each cost of plasticity. Average time values for each set are indicated as black dots within boxplots. In these panels, light blue = *+* and purple = *e*. The total number of runs executed for each different cost of plasticity was 100.

As for frequency data, because higher-cost categories exhibited complete separation, logistic regression could not be estimated beyond the 0% → 1% contrast. Therefore, to support the existence of a tipping point between these two cost values, I applied a binary logistic model comparing only these two levels, which yielded a highly significant reduction in fixation probability (*OR* = 0.05, 95% *CI* = 0.02–0.1, *p* < 0.001). This contrast represents the only statistically identifiable and biologically meaningful transition in the dataset and supports a marked decline in the probability of success of the wild allele when introducing even a low cost of plasticity into the system.

To demonstrate the presence of a further, delayed transition in times to fixation of the mutant allele, given the non-normal distribution of these data, I used a non-parametric approach, i.e., Kruskal-Wallis test to identify global differences across data, and Dunn’s post-hoc to detect the turning point at the proposed threshold. Times differed significantly across plasticity-cost conditions (Kruskal– Wallis test, *p* = 2.29 × 10^-24^). Dunn’s post-hoc test revealed a highly significant difference between 1% and 2% cost conditions (*z* = 5.66, *p* = 1.5 × 10^-8^), with fixation times dropping from a median of 263 to 151 generations. This sharp change indicates that the transition between 1% and 2% cost corresponds to a marked shift in fixation dynamics.

This SA strongly suggests that the inherent cost of plasticity is a pivotal factor affecting the system’s dynamics radically. Importantly, this outcome aligns with general predictions provided by many theoretical models: if the ability to be plastic carries with it some kind of constraints or costs, then the evolution of plasticity is likely to be impeded (Auld et al., 2009; Hendry, 2015). While the convergence between established theory and computational results further corroborates the reliability of *PhePlastiComp* at a system level, these findings also enable the identification of a clear hierarchy between the key modulators of plasticity included in the study. Indeed, 1% cost of plasticity is enough to make the benefits provided by the combined effect of a relatively high degree of plasticity and longer timescales of environmental fluctuations irrelevant in this mixed-strategy system.

This conclusion fits into the broader context of simulation research indicating that inherent costs of plasticity might undermine the potential benefits associated with this strategy, e.g., Miras (2024), that specifically focuses on genetic costs. Since the intrinsic cost of plasticity is explicitly modelled as a separate parameter in *PhePlastiComp* – which allows its isolation from other costs implicit in the system, e.g., the cost related to phenotype-environment mismatching (Auld et al., 2009) – this SA shows that the relevant attenuation of multiple benefits that would normally favour plastic *P. pacificus* genotypes in fluctuating environments is indeed caused by inherent costs.

In addition, the decoupled decays in frequencies and times to fixation along the cost gradient may provide an approximated quantification of threshold values which could apply to the PS312 *P. pacificus* strain and, more broadly, suggest the existence of multi-step, non-linear patterns characterising competition dynamics in this species and comparable organisms displaying bet-hedging and plasticity (see Subsection 3.6 on model applicability). Specifically, the delayed response of the system to increasing costs of plasticity in terms of fixation times of the mutant allele suggests a transitory, yet not negligible coexistence between plastic and non-plastic strategists even when an inherent cost of plasticity (≤ 1%) is present.

### 3.3 The cost of plasticity may induce a switch-like system behaviour and non-plastic genotypes can invade a plastic population across a variety of conditions

The evolutionary dynamics of digital populations of *P. pacificus* were also investigated using more realistic initial conditions, that is, through Invasibility Analyses (IAs). Invasibility is the chance or the ability of a population (in this case, a particular strain/genotype) to increase its frequency from rarity and become established (Pande et al., 2022; Kalirad & Sommer, 2024). All IAs presented here involve a single initial organism belonging to a different strain from the rest of the population, e.g., due to mutation (i.e., *++* → *+e* or *ee* → *+e*) or gene flow. Note that the genotype/strain *+e* is still able to express mouth dimorphism and plasticity but was used as a potential parent of non-plastic worms in scenarios where the prevalent strain was *++*. All following scenarios include 1000 runs.

Although it is widely recognised that constant environments should favour specialist genotypes over generalists (Kassen, 2002; Herron & Doebeli, 2011), the benefits of the former strategy in such environmental conditions have rarely been documented (Condon et al., 2013). The first IA, named “Impact of cost of plasticity on the fixation on non-plastic strains”, provides several insights into this and other related dynamics by monitoring the evolutionary fate of hypothetical *eud-1* mutants/immigrants of *P. pacificus* invading a plastic population.

This scenario is based on the following initial settings: *Initial-++* = 299, *Initial-+e* = 1, *Carrying-Capacity* = 300, stable presence of abundant bacteria, absence of *C. elegans* larvae (= constant environment), *Probability-of-Non-Disjunction-%* = 1, *Degree-of-Plasticity-%* = 10, *Cost-of-Plasticity-%* = 0, 1, 2, and 3, *Cost-of-Starvation-%* = 3, equal benefits in fecundity for both mouth phenotypes = 2 (i.e., *Benefit-of-Faster-Development-%* = *Benefit-of-Predation-%* = 2), and *Efficiency-of-Attack-%* = 8. This scenario aimed at identifying the frequency and time to fixation of a newly arisen allele in response to variation in the cost associated with plasticity.

When *Cost-of-Plasticity-%* = 0, *ƒ_fix_(e)* = 1.5%. This frequency value was more than doubled (3.1%) when *Cost-of-Plasticity-%* = 1, which suggests that the existence of a plasticity-related cost may be critical in increasing the frequency of fixation of the mutant allele. Nevertheless, further increases in the cost of plasticity may have more moderate or even null effects on the capacity of fixation of *e*, that is suggested by outcomes obtained with *Cost-of-Plasticity-%* = 2 (*ƒ_fix_(e)* = 3.9%) and 3 (*ƒ_fix_(e)* = 3.8%) (Figure 6A). Similarly, the inherent cost of plasticity also had a relevant effect on times to fixation, particularly when transitioning from a cost-free condition to *Cost-of-Plasticity-%* = 1, whereas any further increase had more moderate, gradual effects (Figure 6B).

**Figure 6.**
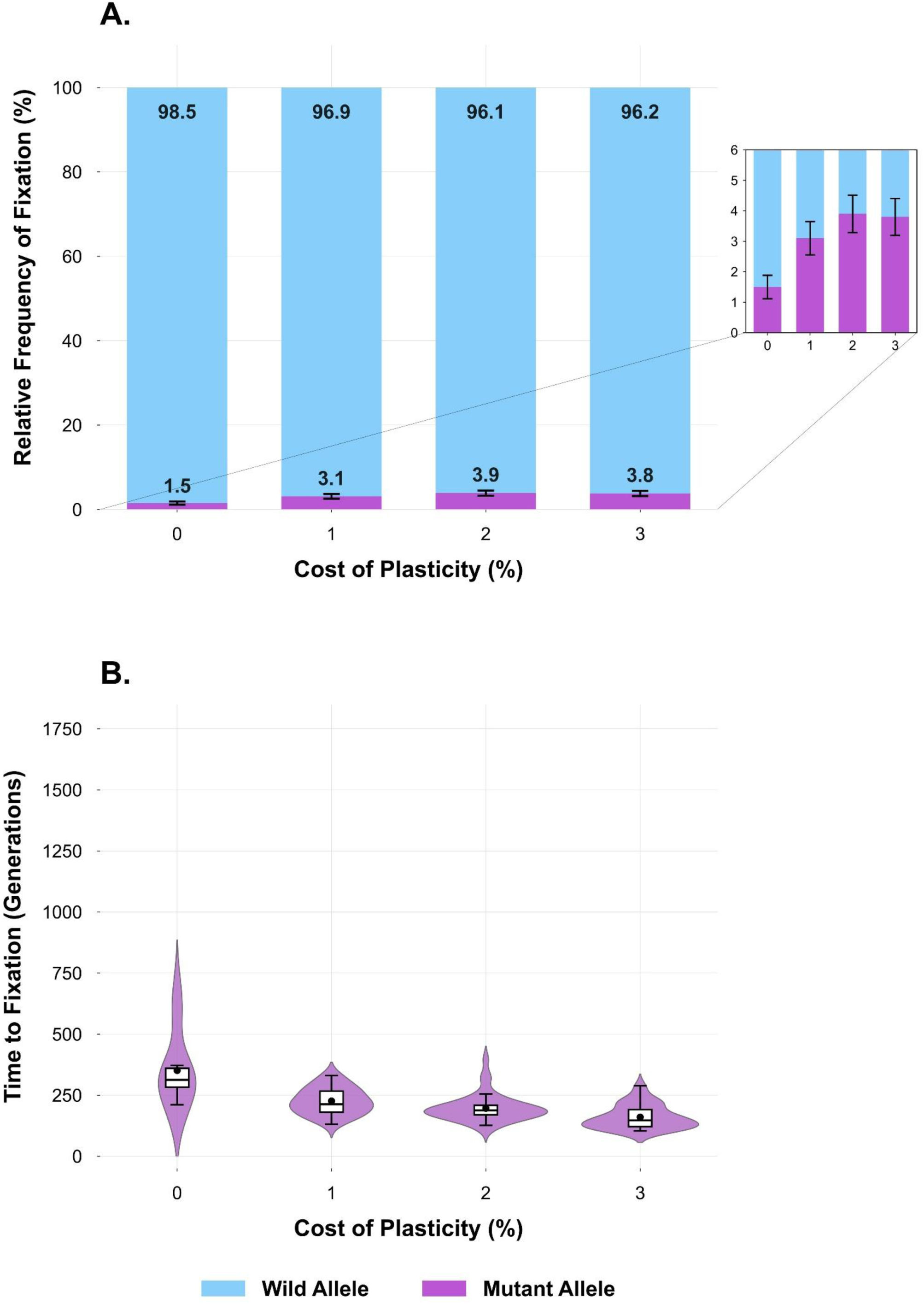
Effects of increasing costs of plasticity on the relative frequencies of fixation of the two competing alleles and times to fixation of the mutant allele. These plots were constructed using data generated by the IA named “Impact of cost of plasticity on the fixation of non-plastic strains”. Panel **A** shows relative frequencies of fixation using stacked bars. Numbers displayed within or above segments indicate the percentage of fixation of each allele, while error bars represent the standard error (SE) of binomial proportions (see also the enlarged detail on the right). Panel **B** shows times to fixation in terms of non-overlapping generations using violin plots with overlaid boxplots to visualise distributions of data across different costs of plasticity. Black dots within boxplots represent mean times to fixation. In these panels, light blue = *+* and purple = *e*. The total number of runs executed for each alternative cost of plasticity was 1000.

To quantify how *ƒ_fix_(e)* changed across sequential increments in the cost of plasticity, I fitted three separate logistic regression models, each using a different cost as the reference category. Frequency of fixation increased sharply from 0% to 1% (*OR* = 2.1, 95% *CI*: 1.13–3.92, *p* = 0.02), which supports the existence of a clear phase transition. The increase from 1% to 2% was modest and statistically non-significant (*OR* = 1.27, 95% *CI*: 0.78–2.05, *p* = 0.33). Finally, *ƒ_fix_(e)* showed no appreciable difference between 2% and 3% (*OR* = 0.97, 95% *CI*: 0.62–1.53, *p* = 0.91), consistent with a plateau at higher costs.

To assess whether time to fixation data would also confirm the presence of a phase transition between 0% and 1% cost of plasticity, I first used a global Kruskal–Wallis test, that revealed significant differences in times until fixation across alternative costs (*p* = 4.42 × 10^-9^), then Dunn’s post-hoc on the 0% → 1% contrast, obtaining again a significant p-value (*z* = 2.15, *p* = 0.03), with median fixation times dropping from 313 to 213 generations.

Unexpectedly, these outcomes inform that, while the existence of an intrinsic cost of plasticity may uncover a switch-like behaviour of the system by rapidly increasing the probability of success of a newly arisen allele reducing environmental sensitivity, advantages provided to this allele do not steadily increase as the cost of plasticity increases beyond a certain threshold. A plausible explanation for the emergence of this constrained steady state is that the inherent cost of plasticity must also be shouldered by heterozygotes (i.e., invaders) carrying a single mutated *eud-1* copy, since they are still plastic. This may impose an upper limit to the chances of invasion by non-plastic mutants which, although capable of producing cost-exempted offspring, must face a “head start” penalty in addition to their scarce initial frequency.

To ascertain whether this complex pattern generated in response to increasing costs of plasticity – i.e., a relevant increase in the frequency of fixation of the mutant allele followed by a stabilisation phase – may be regarded as a robust property of the modelled system, I also carried out a sensitivity analysis using this IA as a baseline. More generally, this analysis aimed at testing the robustness of the ABM in response to variation in parameter values which I always kept fixed in all other scenarios, namely *Carrying-Capacity* and *Cost-of-Starvation-%*. To do this, I compared values of *ƒ_fix_(e)* obtained using the default settings of this IA (including *Carrying-Capacity* = 300 and *Cost-of-Starvation-%* = 3) with values of *ƒ_fix_(e)* obtained using *Carrying-Capacity* = 100 and *Cost-of-Starvation-%* = 1, respectively (everything else being equal). This exploration of the parameter space was conducted for each set of runs identified by an alternative value of *Cost-of-Plasticity-%* (0, 1, 2, and 3).

The comparison between frequency data across different costs of plasticity revealed that the model is generally robust and that the trends suggested by its outcomes may be informative for population sizes ≤ 300 and moderate-to-low costs associated with bacterial starvation (Figure 7), although it is important to notice that the tipping point in terms of benefits conferred to the mutant allele might arise with a slightly higher value of *Cost-of-Plasticity-%* (i.e., 2 instead of 1) when using a very small population of virtual nematodes, i.e., 100 individuals. In addition to being possibly found in *P. pacificus*, I speculate that the non-linear pattern detected here could also characterise other comparable dimorphic taxa where polyphenism is regulated upstream by dosage-dependent switch genes, as the maintenance of the ability to develop both phenotypes by heterozygotic genotypes would likely imply a retention of molecular costs inherent to the decision-making process (see also Subsection 2.4).

**Figure 7.**
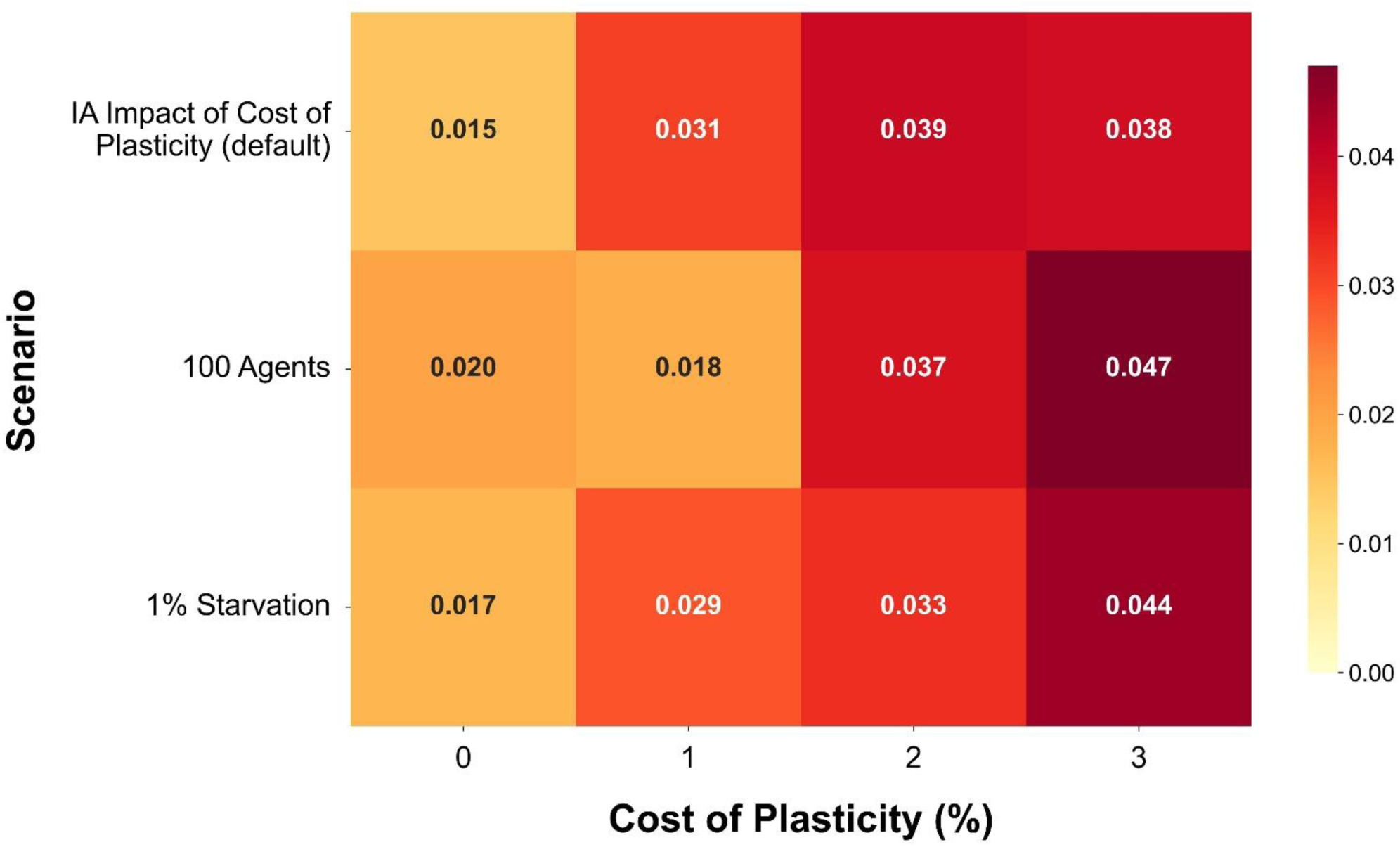
Heat map of frequencies of fixation of the mutant allele in response to variation in the inherent cost of plasticity across alternative settings. This sensitivity analysis was conducted using alternative versions of the IA scenario “Impact of cost of plasticity on the fixation of non-plastic strains”. Darker shades indicate higher relative frequencies of fixation of the mutant allele, whereas lighter shades indicate lower frequencies. Frequency values obtained under each condition are also reported within each cell. The total number of runs executed for each tested condition was 1000.

Given the relevance of this scenario for the purposes of the study, an additional version of this IA was carried out using the same basic settings of “Impact of cost of plasticity on the fixation of non-plastic strains”, this time with fixed *Cost-of-Plasticity-%* = 1, an environment fluctuating at an intermediate rate (*Environmental-Cycles* = 10), and *Degree-of-Plasticity-%* = 20, which would be expected to favour the plastic strain. Surprisingly, the non-plastic strain still reached fixation in 7/1000 replications (0.7%), with *t̅_fix_(e)* = 538.14 (IQR = 171, Q1 = 423.5, Q2 = 488, Q3 = 594.5). Therefore, non-plastic strains can invade plastic populations even when the dietary conditions fluctuate over time.

This finding reveals that the mutant allele reaches fixation not only when the amount of available microbial food is abundant and stable over time, but even when the environment is periodically oscillating in terms of food sources and non-plastic strains have to face multiple time intervals in which they must pay a cost due to bacterial starvation (Serobyan et al., 2014), as long as plasticity has at least a minimum cost (i.e., 1%). I argue that this counterintuitive observation could be explained by considering that, although non-plastic strains pay a starvation-related cost, this cost is also shared with plastic strains, that must additionally bear the cost of plasticity. Furthermore, differently from the cost of starvation, the intrinsic cost of plasticity is not cyclic. This means that while non-plastic strains can recover from diminishing densities during phases when microbial food is abundant (on moderate timescales of dietary fluctuation), the benefits gained by expressing higher percentages of the optimal morph in the proper environment might not be sufficient for the maintenance of a high net advantage by plastic strains.

Therefore, while supporting the benefits of developing a single, optimal phenotype in a static environment, simulations also suggest the possible advantages of unconditional expression of a single phenotype in a fluctuating environment, assuming that plasticity is costly *per se*. In other terms, regardless of environmental variability, the existence of an intrinsic (even if low) cost of plasticity seems to be a sufficient condition to guarantee chances of invasion to strains defective in the expression of the EU morph.

This surprising phenomenon has already been reported in a bacterial model. In particular, the glycolysis pathway can switch from conditional to constitutive expression when colonies experience a regularly fluctuating environment over time, even if an inherent cost of plasticity is absent (Herron & Doebeli, 2011). However, since most natural isolates of *P. pacificus* are dimorphic and EU-biased (Ragsdale et al., 2013), it could be hypothesised that invasions by *eud-1* mutants are unlikely due to the absence of an intrinsic cost of plasticity, presence of fractional costs < 1%, or reduction in fitness experienced by some mutants (e.g., caused by pleiotropy, that was not explicitly modelled in *PhePlastiComp*, see Subsection 3.5).

### 3.4 Invasions by plastic genotypes might be highly dependent on the cost of plasticity and difference in fitness between phenotypes

While all previous scenarios included equal benefits in fecundity for both mouth forms (i.e., ST and EU), the IA “Impact of differences between the two benefits” was designed to assess the evolutionary effects of differences between these two benefits (henceforth Δ*b*_EU-ST_, i.e., *Benefit-of-Predation-%* - *Benefit-of-Faster-Development-%*). In fact, it is reasonable to hypothesise that differences in fecundity between ST and EU nematodes may favour or harm each strain in relation to its ability to express the phenotype most attuned to the current environmental conditions with appreciable frequencies. Moreover, since the ST morph is shared by both strains, whereas the EU phenotype is only developed by carriers of at least one copy of the wild-type allele, the global dynamics resulting from bringing imbalances between the two benefits could uncover several unpredictable patterns.

This scenario is based on the following initial settings: *Initial-ee* = 299, *Initial-+e* = 0 or 1, *Initial-++* = 0 or 1, *Carrying-Capacity* = 300, alternate presence of abundant bacteria and *C. elegans* larvae (= fluctuating environment), *Environmental-Cycles* = 10, *Probability-of-Non-Disjunction-%* = 1, *Degree-of-Plasticity-%* = 0 or 20, *Cost-of-Plasticity-%* = 0 or 1, *Cost-of-Starvation-%* = 3, *Benefit-of-Faster-Development-%* = 1, *Benefit-of-Predation-%* = 1, 2, or 3, and *Efficiency-of-Attack-%* = 8. This specific timescale of environmental fluctuations and level of plasticity were chosen because intermediate environmental cycles and higher developmental sensitivity might be advantageous for plastic strategists (see Subsection 3.1).

When the genotype of the initial invader was *+e* and *Cost-of-Plasticity-%* = 1, the outcomes were perhaps surprising: in all three sets (i.e., when Δ*b*_EU-ST_ = 0, 1, and 2, respectively), the wild-type allele never reached fixation in any of the 1000 runs carried out for each set, indicating that advantageous parameter values for plastic strains combined with remarkable benefits conferred to the EU phenotype were not enough to alleviate the effects of a low cost of plasticity. When the genotype of the initial invader was *++* the outcomes were even more unexpected, since this genotype has a 100% probability to develop the advantaged morph (EU) when bacteria are scarce (*Degree-of-Plasticity-%* = 20). When Δ*b*_EU-ST_ = 0% or 1%, the *+* allele was still not able to reach fixation in any of the 1000 replications. A 2% difference in favour of the EU phenotype was critical to enable a very low number of fixations (3/1000).

These initial results inform that the presence of a minimum integer cost associated with the expression of plastic responses might be an insurmountable barrier for mutants reverting to a plastic genotype, as well as for single plastic immigrants invading a non-plastic population. Consequently, more structured insights may only be provided by considering an alternative context denoted by intrinsic costs of plasticity < 1%, i.e., either fractional or null costs.

An alternative starting condition involving a *+e* invader, this time with exclusive stochastic regulation of the plastic response (*Degree-of-Plasticity-%* = 0) and everything else being equal, revealed that a benefit in fecundity for EU higher than the benefit associated with ST may lead to several fixations of the wild-type allele, i.e., in 9 runs out of 1000 (*ƒ_fix_(+)* = 0.9%). Although a single fixation event was also observed with equal benefits in fecundity for both phenotypes, any asymmetry favouring EU seems to be an essential requirement to ensure several fixations of *+* (Figure 8A), which could indicate a further non-linear behaviour of the modelled system. Despite this first indication, I also tested whether increasing the level of plasticity up to 20% would further enhance the frequency of fixation of the wild-type allele under the same settings and observed a slightly increased advantage for *+* when Δ*b*_EU-ST_ = 2%, as fixation was reached in 13 runs out of 1000 (*ƒ_fix_(+)* = 1.3%). Conversely, fixation was reached only in 1 and 6 runs out of 1000 (*ƒ_fix_(+)* = 0.1% and *ƒ_fix_(+)* = 0.6%) when Δ*b*_EU-ST_ = 0% and 1%, respectively (Figure 8B).

**Figure 8.**
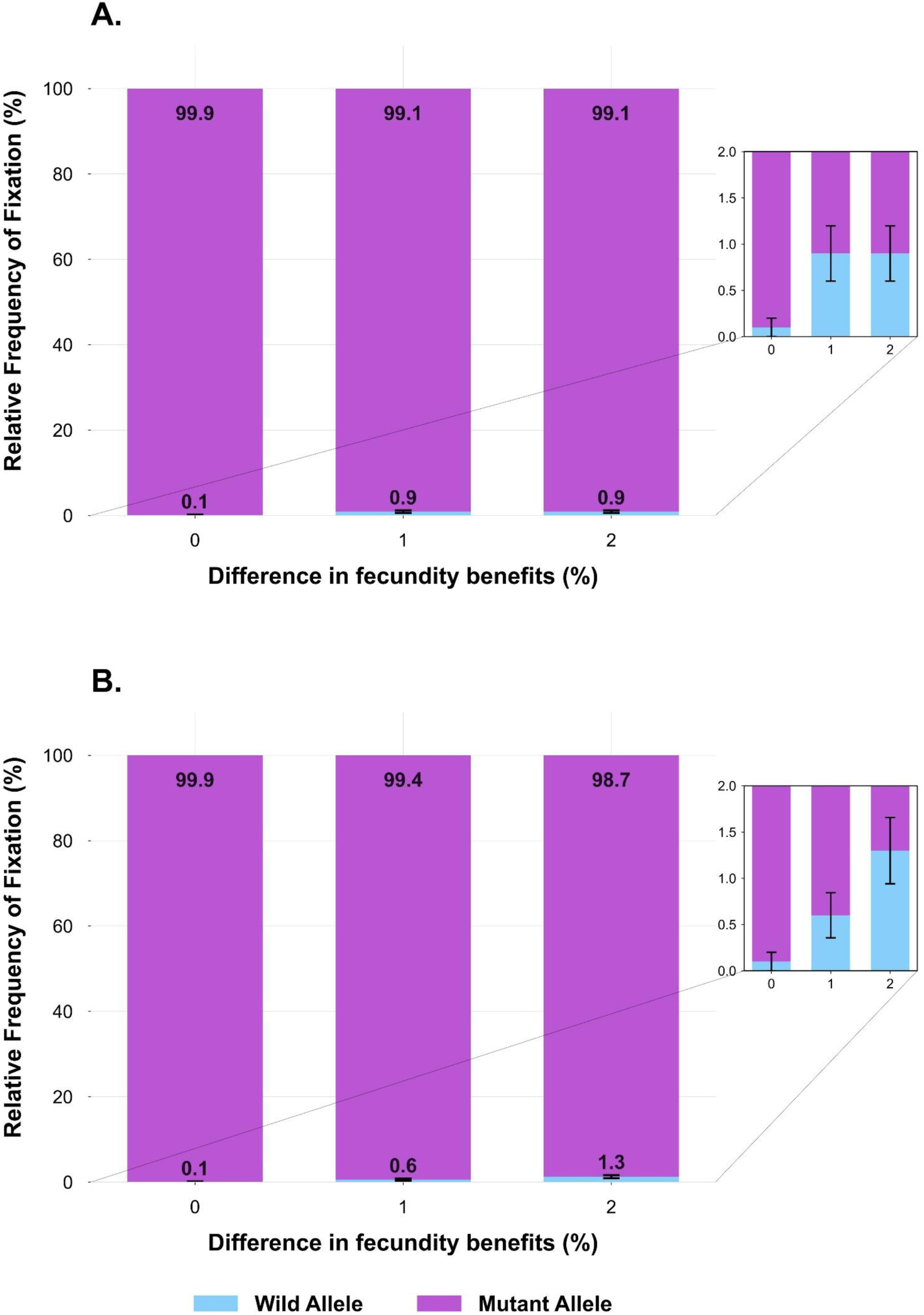
Effects of increasing differences between the two benefits in fecundity on the relative frequencies of fixation of the wild-type allele. These plots were constructed using data generated by the IA named “Impact of differences between the two benefits”, assuming either the absence of inherent costs of plasticity or the presence of irrelevant fractional costs. Relative frequencies of fixation of the two competing alleles are represented by stacked bars. Numbers displayed within or above segments indicate the percentage of fixation of each allele, while error bars indicate the standard error (SE) of binomial proportions (see also the enlarged details on the right). Panel **A** shows a version of the IA denoted by plastic genotypes which are exclusive or nearly exclusive bet-hedgers, i.e., with *Degree-of-Plasticity-%* = 0. Panel **B** shows an alternative version of the IA where plastic genotypes have the maximum possible level of plasticity selectable in the model (*Degree-of-Plasticity-%* = 20). In these panels, light blue = *+* and purple = *e*. The total number of runs executed for each alternative difference in fecundity benefits was 1000.

Therefore, it seems that these two alternative strategies adopted by plastic genotypes, namely bet-hedging and bet-hedging + conditional plasticity, may only be effective when the benefits in fitness experienced by the predatory phenotype are higher than those experienced by the bacterivorous morph, and only under cost-free or nearly-cost-free conditions. In addition, the relationship between differences in fecundity benefits and frequency of fixation of the wild allele seems to be switch-like for exclusive bet-hedgers and linear for strains showing a response to the environment.

To provide formal support to the latter observations, I first quantified how *ƒ_fix_(+)* responded to increasing differences in fitness (0%, 1%, 2%) using stepwise logistic regressions, each with a different reference level, in the IA version including exclusive bet-hedgers. *ƒ_fix_(+)* showed a sharp phase transition between 0% and 1% (*OR* = 9.07, 95% *CI*: 1.15–71.74, *p* = 0.04), indicating that even a 1% fitness advantage experienced by EU nematodes strongly promotes fixation of *+*. In contrast, the change from 1% to 2% yielded an *OR* = 1 (95% *CI*: 0.39–2.53, *p* = 1), demonstrating a clear plateau where additional fitness/fecundity differences no longer increase *ƒ_fix_(+)*.

Conversely, the linear term produced by a logistic regression performed on data generated from the IA version including bet-hedging + conditional plasticity showed an *OR* = 2.91 (95% *CI*:1.45-5.81, *p* = 2.51 × 10^-3^), clearly demonstrating that increasing the difference in fecundity between the two morphs provides linear increases in the advantages conferred to plastic genotypes.

A plausible mechanistic explanation for these patterns is that higher benefits attributed to the phenotype that is only developed by plastic strains (EU) may greatly enhance the reproductive performance of these organisms when prey larvae are present, whereas non-plastic strains (which always express the ST morph) are only slightly advantaged during cycles when microbial food is abundant. Moreover, the only explicit cost in fecundity shared by both strategies (i.e., the cost of starvation), is experienced by all nematodes for the same time intervals.

Collectively, these findings suggest that any inherent cost of plasticity ≥ 1% would act as a shield for a population expressing the ST phenotype unconditionally and may have a more relevant impact than the combined effect of an intermediate timescale of environmental fluctuation and a relatively high level of plasticity (20%), which tended to favour plastic strains in SAs. While confirming the hierarchy of evolutionary modulators outlined in Subsection 3.2, the observation that fitness unbalances favouring the only exclusive phenotype developed by plastic genotypes might enable fixation of such genotypes – with alternative patterns depending on the degree plasticity – supports the relevance of this additional component in directing competition dynamics.

However, as this positive outcome for plastic strains was only observed when using a null/irrelevant intrinsic cost of plasticity, the latter factor may still be regarded as the most prominent modulator of competition in this mixed-strategy system. This analysis clearly shows that the pressure exerted by costs in terms of channelling evolutionary trajectories is more effective than the synergistic advantages provided by greater levels of plasticity, intermediate environmental cycles, and appropriate allocation of phenotype-specific benefits.

Since wild isolates of *P. pacificus* regularly show both stochastically determined, strain-specific phenotype ratios and conditional regulation of plasticity (Lenuzzi et al., 2021), this conclusion prompts a deeper reflection on the nature and management of costs of plasticity in this species. Potential explanations include even higher fitness benefits experienced by EU worms, influence of the spatial component on population dynamics, absence of intrinsic costs of plasticity, and presence of fractional costs < 1%. While very high EU-related benefits could theoretically account for the observed spread of plastic strains, the biological plausibility of such a pronounced fitness gap between morphs (> 2%) remains to be established, as it could confer unrealistic selective advantages to non-plastic predatory strains (i.e., full-EU). In contrast, alternative modelling frameworks have already demonstrated that spatial and temporal heterogeneity may alleviate the cost of plasticity in this species (Kalirad & Sommer, 2024A) (see also Subsection 3.5).

I argue that the explanatory power of spatial resource distribution is further enhanced when combined with the intrinsic logic underlying plastic strategies evolved by *P. pacificus*. Specifically, the inherent cost of plasticity might have been largely circumvented by evolving a highly asymmetrical mixed strategy which is closer to pure bet-hedging than to environmental responsiveness, so that the actual cost that individuals must bear could mostly be due to phenotype-environment mismatching (Auld et al., 2009). This would perfectly fit with developmental patterns observed in wild isolates, in which the degree of developmental sensitivity is much lower than that shown by other diplogastrid nematodes not belonging to the genus *Pristionchus* (Susoy et al., 2015). This speculation is also supported by evidence suggesting that the fitness cost related to additional regulatory machinery may be inappreciable in most cases (Latta et al., 2012; Murren et al., 2015), and that an inherent cost of plasticity is mainly found and expected in highly plastic populations (Lind & Johansson, 2009).

Notably, this does not necessarily rule out the existence of inherent costs, as the same reasoning would apply in the event of costs lower than the 1% threshold detected in this study. However, as the current version of *PhePlastiComp* was designed to only include integer fitness costs to focus on more macroscopic transitions, a more precise identification of the threshold above which the evolution and maintenance of plasticity is unlikely in *P. pacificus* would need to be addressed in future, fine-grained assessments of this critical variable.

To sum up, the interplay between degree of plasticity, cost of plasticity, and difference in fitness between phenotypes proves to be crucial in this dimorphic species and deserves further investigation aimed at explaining how a plastic genotype could prevail in environments changing over time that are dominated by non-plastic strategists.

### 3.5 Model limitations

Although *PhePlastiComp* provides a suitable simulation platform for exploring eco-evolutionary dynamics combining stochastic, conditional, and constitutive phenotype expression, its predictive scope is defined by specific design choices.

First, considering the ecological niches where *P. pacificus* naturally interacts, the assumption of a well-mixed environment is a deliberate simplification. Recent studies also focusing on *P. pacificus* have already explored more realistic environmental conditions, e.g., Kalirad & Sommer (2024A), that implements a stage-structured model demonstrating how spatial and temporal heterogeneity in the distribution of resources in a metapopulation can indeed affect the outcomes of strain competition and mitigate the cost of plasticity. However, my model focuses on a different theoretical objective: isolating the intrinsic selective effects of key variables without the confounding influence of spatial dynamics (e.g., due to dispersal and refugia). This abstraction provides a rigorous baseline framework to assess parameter interrelationships and trade-offs independently from space-related mechanisms, which are however among the most relevant possible inclusions in future versions of the ABM.

Second, the model excludes *de novo* mutations occurring during runs. While this could also be an additional element of realism to be considered for a future update, this exclusion could currently limit the exploration of potential deviations in the evolutionary trajectories, at least in scenarios occasionally showing longer coexistence between strains. For instance, this might apply to some IAs, where new mutants arisen from the original population could provide a limited boost to the probability of fixation of the most infrequent strategy when it is still growing in number.

Third, the fixed male production rate = 1%, while anchored on empirical data and highly characterising *P. pacificus*, is likely not critical in directing the global competition dynamics, as it introduces a negligible percentage of cross-fertilisation events for each generation. Consequently, while the simulated trends may also apply to organisms with 100% clonal reproduction, the inclusion of this variable into the model might even enable future investigation aimed at identifying thresholds beyond which the model could approximate a dioecious system.

Fourth, the model focuses on any hypothetical mutation exclusively affecting mouth-form ratios, which excludes pleiotropic effects affecting non-plastic genotypes. While this would limit potential comparisons of the model’s outcomes with assays involving mutagenised *eud-1* nematodes – which often exhibit additional morphological and locomotory side effects reducing their fitness – this choice enables possible comparisons involving full-ST natural isolates. Interestingly, by applying a difference in benefits that largely favours plastic, *++* nematodes (as they are 100% EU under bacterial starvation), the IA “Impact of differences between the two benefits” could indeed approximate a context involving pleiotropy-related costs for non-plastic strains.

### 3.6 Model applicability

Notwithstanding the above limitations, I argue that the outcomes of *PhePlastiComp* – with the necessary caution due to the specific parameterisation – could also suggest qualitative patterns potentially detectable in other taxa sharing fundamental structural similarities with *P. pacificus*. Indeed, while life history characteristics are specifically tailored to the model organism, the trends uncovered in this study could apply to organisms possessing the following attributes:

- High prevalence of self-fertilisation or asexual reproduction;
- Stochastically and conditionally regulated dimorphism;
- Dimorphism enabled by a developmental switch;
- Unbalanced phenotype ratios in plastic genotypes.

Importantly, since generations are non-overlapping and environmental cycles were designed to match the duration of the former, there are no specific constraints related to *P. pacificus* life cycle which would preclude the extrapolation of these findings to species with substantially different lifespans. Thus, beyond nematodes, potential candidates for this framework might be found among diverse taxa including arthropods and plants (Grantham et al., 2016; Van Den Elzen & Emery, 2025).

### 3.7 Synergistic strategies to cope with climate change

Eco-evolutionary trade-offs between bet-hedging, conditionally regulated plasticity, and environmental homeostasis transcend the specific field of application and relate this study and its main findings to the critical issue of climate change in a deep way. Indeed, understanding the strategies through which organisms respond to changing environments is crucial to forecast extinction risks of natural populations (Vinton et al., 2022).

As climate change may consist of a combination of incremental shifts and intensifying extremes, an assessment of the matching between alternative strategies and specific environmental conditions is critical. In this regard, a recent agent-based model focusing on plants and strategies adopted by farmers – adjusting calendars, switching crops, diversifying production – has addressed this key issue by detecting the most suitable strategy under different conditions: plasticity was found to be most advantageous under gradual shifts, whereas bet-hedging buffered volatility in unpredictably fluctuating conditions (Favretto et al., 2026). Notably, the authors highlighted that the distinction between these two strategies blurs in practice, suggesting that plasticity and bet-hedging are often part of a continuum rather than mutually exclusive alternatives.

The outcomes of *PhePlastiComp* could successfully expand this perspective by indicating that asymmetric mixed-strategies involving prevalent bet-hedging and relatively low levels of plasticity – adopted by taxa reflecting the conditions listed in Subsection 3.6 – could be more efficient than fixed phenotype production across a range of temporal fluctuation rates (fast, moderate, and slow), provided that plasticity has very low or null intrinsic costs. Moreover, the protective effect of this complex strategy would be amplified if the phenotype that can only be developed by plastic strategists has higher fitness benefits than the shared phenotype. Nevertheless, very rapidly oscillating environments or unpredictable patterns of change could still reduce or even eliminate these advantages, making combined strategies far from being universally applicable solutions.

## 4. Conclusion

The opportunity to model the complex interplay between environment, development and genetics, combined with a focus on individual heterogeneity, makes agent-based modelling an ideal approach to generate theoretical insights by relying on empirical information from species that have already proven to be powerful experimental subjects.

While proposing *PhePlastiComp* as a reliable simulation platform aimed at investigating competition dynamics in *P. pacificus*, this exploratory analysis provides novel hypotheses and unprecedented details which shed light on the complex interplay between plastic, stochastic, and constitutive strategies in this nematode species and comparable dimorphic systems.

## Acknowledgements

I wish to thank Professor Ralf J. Sommer and Dr. Ata Kalirad of the Max Planck Institute for Biology of Tübingen (Integrative Evolutionary Biology) for their valuable suggestions and helpful discussion while developing the chapter of my PhD thesis on which this article is based. I also wish to thank my PhD thesis advisors Gianluigi Oliveri, Valentino Romano and Marco Carapezza of University of Palermo for their support during my doctoral studies and Paco Garcia-Gonzalez of Doñana Biological Station (EBD CSIC Seville) for all the experience gained working with him on similar research projects.

## Supplementary Material

**Supplementary File 1. *PhePlastiComp*: Overview, Design Concepts, and Details (ODD).** ODD protocol of the NetLogo agent-based model.

**Supplementary File 2. Relevant details about *Pristionchus pacificus* for the implementation of the agent-based model.** Summary of the information extracted from the literature about *P. pacificus* genetics, development, and ecology and used to implement the agent-based model.

**Supplementary File 3. Macro-validation of the model.** Preliminary scenarios aimed at supporting the reliability of *PhePlastiComp* by showing a set of expected, basic emerging patterns.

## Data availability

The NetLogo model I implemented and used for simulating my scenarios can be freely downloaded at the following url: https://doi.org/10.5281/zenodo.20452270. The model can be opened by installing the NetLogo simulation platform (version 6.2.0), which can be downloaded free of charge from the official website of the software at the following link: https://www.netlogo.org/downloads/archive/6.2.0/. Data extracted from NetLogo runs and used for the analysis of the model can also be freely downloaded at the following url: https://doi.org/10.5281/zenodo.20452111.

## Declaration of generative AI and AI-assisted technologies

During the preparation of this work I used Gemini in order to refine the Python scripts used to generate Figures 3-8 and perform statistical tests, which are completely based on my original NetLogo data. After using this tool/service, I reviewed and edited the content as needed and take full responsibility for the content of the published article.

