## Supplementary File 1 for "Agent-based modelling of a nematode system provides novel insights into the evolutionary constraints and modulators of phenotypic plasticity, bet-hedging, and environmental homeostasis"

*PhePlastiComp*: Overview, Design Concepts, and Details (ODD)

### 1. Overview

#### 1.1 Purpose and patterns

The purpose of the model is to simulate the competition between plastic and non-plastic genotypes (depending on the state of the developmental switch gene *eud-1*) of the dimorphic nematode worm *Pristionchus pacificus* over multiple, non-overlapping generations, in an in vitro context with periodic environmental fluctuations. More specifically, the model was conceived to investigate the impact of the interplay between an intrinsic cost of plasticity, the timescale of environmental fluctuations, and the individual degree of plasticity on the evolutionary trajectories of populations of *P. pacificus*, which could also provide some more general suggestions on other comparable systems. Here I define i) the *intrinsic cost of plasticity* as a reduction in the number of offspring produced by an individual as a consequence of its ability to develop alternative phenotypes depending on environmental stimuli, ii) the *timescale of environmental fluctuations* as a time interval, expressed in terms of non-overlapping generations, in which one food source (out of two possible sources) prevails over the other before the ratio between the two sources is reversed, and iii) the *individual degree of plasticity* as the magnitude of the phenotypic response to the environmental condition experienced (i.e., the probability that an individual develops a specific phenotype following exposure to an environmental trigger).

The main objective of simulations is to generate data on the frequency and time to fixation/extinction of the two competing genotypes and their related alleles (i.e., variants of *eud-1*). Indeed, the patterns of fixation/extinction of genetic variants are influenced by all genetic, environmental, and developmental factors that can affect the fitness of the phenotypes that such variants contribute to producing, which makes the model suitable for obtaining information on the relationship between these factors and population-level evolution.

#### 1.2 Entities, state variables, and scales

The individual entities/agents included in the model are nematodes belonging to two different species, i.e., *P. pacificus* and *Caenorhabditis elegans*, *Escherichia coli* bacteria, and NetLogo patches (grid cells). The NetLogo observer is the only system-level agent.

*P. pacificus*, which is the main type of agent, is an established model organism in the field of evolutionary developmental biology (Schroeder, 2021), and here is used to study the competition between plastic and non-plastic genotypes/strains/strategies. *C. elegans* is a potential prey of *P. pacificus*, whereas *E. coli* is the main food source of the latter (Serobyan et al., 2014). The set of all patches constitutes the in vitro environment where all these organisms are located, e.g., a Petri dish. The observer is used to give instructions to all other classes of agents.

General information on *P. pacificus* genetics, development, ecology, morphology, and eating behaviour is provided in Supplementary File 2 (“Relevant details about *Pristionchus pacificus* for the implementation of the agent-based model”). That document is both a rationale, as it justifies the choice of entities included in the model, and a summary of the main characteristics of *P. pacificus* that were implemented in *PhePlastiComp*. Together with this ODD protocol, the appendix is also useful from a micro-validation perspective (Wilensky & Rand, 2015), as it provides the observational and experimental data drawn from the literature based on which the agents were programmed.

All state variables associated with individual and system-level agents, i.e., variables that change over simulation time or across agents, are listed in Table S1.

**Table S1. State variables included in the model.** The table summarises all state variables included in the model, grouped by agent class and accompanied by the type of variable, the range of all possible values for that variable, and a short description. The value of dynamic variables changes over simulation time, whereas the value of static variables does not.

| **Agent class** | **State variable** | **Type** | **Range** | **Description** |
| --- | --- | --- | --- | --- |
| Observer (system-level agent) | *stored-cost-of-starvation-%* | Integer number | 0-5 | Static variable. Memory of the cost in fecundity due to bacterial starvation in time intervals in which bacterial food is abundant |
| All individual agents | *who* | Integer number | Any value | Static variable. Unique ID number of each agent |
| All individual agents | *color/pcolor* | Integer number | - | Static variable. NetLogo built-in primitive indicating the colour of each agent |
| All nematodes and bacteria | *heading* | Integer number | 0-360 | Dynamic variable. NetLogo built-in primitive indicating the orientation angle of moving agents |
| All nematodes and bacteria | *xcor*, *ycor* | Real number | -32-32 | Dynamic variable. NetLogo built-in primitives indicating the coordinates of moving agents |
| All nematodes and bacteria | *shape* | Literal | - | Dynamic variable. NetLogo built-in primitive indicating the shape of moving agents. Bacteria and *C. elegans* worms have fixed shapes, whereas the shape of each *P. pacificus* worm changes during its lifetime (i.e., from egg to juvenile worm, and from juvenile worm to adult worm with one of two alternative mouth phenotypes, ST or EU) |
| All nematodes and bacteria | *size* | Real number | Any value | Dynamic variable. Bacteria and *C. elegans* worms have different sizes. In *P. pacificus*, size of both eggs and juvenile worms increases as developmental stages progress |
| All nematodes | *species* | Literal | - | Static variable. Species of each nematode (i.e., *C. elegans* or *P. pacificus*) |
| *P. pacificus* nematodes | *genotype* | Literal | - | Static variable. Genotype of each nematode. Since the single locus of interest (*eud-1*) is X-linked and non-disjunction events can occur, there are 5 possible genotypes: *++*, *+e*, *ee*, *+O*, and *eO*, where *+* is the wild-type allele and *e* is the mutant allele |
| *P. pacificus* nematodes | *phenotype* | Integer number | 0-2 | Dynamic variable. Phenotype of each nematode. Juvenile worms have phenotype 0 (undifferentiated mouth-form), whereas adult worms can develop either phenotype 1 (EU mouth-form, facultative predators) or phenotype 2 (ST mouth-form, bacterivorous) irreversibly |
| *P. pacificus* nematodes | *sex* | Literal | - | Static variable. Worms can be either hermaphrodites or males |
| *P. pacificus* nematodes | *partner* | Literal/Integer number | - | Dynamic variable. Hermaphrodite worms which reproduce via self-fertilisation have no sexual partner (“nobody”) for all their lifetime, whereas adult hermaphrodites crossing with males have a single partner identified by its unique ID number (*who* number). Male worms also start with no partner and, when they are adult, they have a single partner identified by its unique ID number (*who* number) |
| *P. pacificus* nematodes | *developmental-stage* | Literal | - | Dynamic variable. Nematodes start with no developmental stage (“none”), then they go through the following stages: new-egg, J1, J2, J3, J4, and adult |
| *P. pacificus* nematodes | *developmental-progression* | Integer number | 0-820 | Dynamic variable. Nematodes live for 820 NetLogo time steps (ticks). In the model, 10 ticks = 1 hour |
| *P. pacificus* nematodes | *time-to-oviposition* | Integer number | 0-120 | Dynamic variable. Countdown from the start of adulthood to oviposition. Nematodes lay their eggs simultaneously and the surviving eggs make up the next generation of worms |
| *P. pacificus* nematodes | *generation* | Integer number | Any positive value | Static variable. Generation to which the individual belongs (first generation = 0, second generation = 1, and so on) |
| *P. pacificus* nematodes | *parent-1* | Integer number | - | Dynamic variable. This value is simply the ID number (*who*) of the individual, except for the first time step (tick) of life of newly generated offspring. In the latter case, this value is the ID number of the hermaphrodite parent (which is usually the only parent). Both this variable and *parent-2* have the only function to assign costs in fecundity to each single offspring correctly (i.e., without affecting offspring of other individuals) |
| *P. pacificus* nematodes | *parent-2* | Literal/Integer number | - | Dynamic variable. In the first generation, this value is “nobody” for all juvenile worms and always “nobody” for all male worms. Adult hermaphrodites which cross with a male change this value to the ID (*who*) number of their male partner. In the offspring of hermaphrodites which crossed with males, this value is the ID number of the father and is replaced by the ID number of the male partner if they are hermaphrodites and cross with a male when they are adult |
| *P. pacificus* nematodes | *nematodes-killed* | Integer number | 0-5 | Dynamic variable. Number of *C. elegans* larvae eaten by each worm with *phenotype* = 1 (EU). The upper limit of 5 was chosen so as not to unrealistically increase the benefits of predation, which also depend on this variable |
| Patches (grid cells) | *pxcor*, *pycor* | Integer number | -32-32 | Static variable. NetLogo built-in primitives indicating coordinates of fixed agents (grid cells) |

The model includes a simplified spatial component, as all bacteria and nematodes change their location during runs by moving above a virtual representation of a culture plate. The reason why I implemented a basic in vitro environment instead of a natural environment with more realistic elements and/or barriers is the clear focus of the model on temporal, rather than spatial variation in resource availability. The simulation space is homogeneous and toroidal, and it is represented by a discrete grid composed of square cells (patches), although all organisms move on a continuous space where locations are identified by real numbers. The spatial extent of the model is a square of 35x35 patches, and there is no strict proportional relationship between the size of this space and that of a real culture plate, although it can reasonably be assumed that the area of a patch would correspond to approximately 3-5 mm^2^ of a real plate (I increased the size of moving agents compared to the space they would actually occupy in a real culture plate, especially *P. pacificus* worms, to improve their visualisation in the NetLogo interface).

Time is represented using discrete time steps called ticks, and simulation updates are provided at each tick. In the model, a tick is the time interval required for all agents to execute all the algorithms assigned to them one time (i.e., a single iteration). The order in which each agent executes the programme within a tick is irrelevant in this context, except for the last tick of life of *P. pacificus* worms: in this case, all adults must first generate their offspring, then apply any costs in fecundity, and only then die. 10 time steps/ticks within simulations correspond to 1 hour, and the duration of each non-overlapping generation of *P. pacificus* is 820 time steps/ticks (= 82 hours, that is the approximate lifespan of *P. pacificus* in the real world (Hong & Sommer, 2006)). In other words, there is a specific proportion between simulation time and real time (see also Supplementary File 2 and Subsection 3.3 in this document for further details). Each run can potentially have any duration, both in terms of ticks and generations, but in this study, runs are stopped when an absorbing state (i.e., allele fixation/extinction) is reached.

#### 1.3 Process overview and scheduling

In this subsection, I provide a comprehensive list of all processes executed by the model at each time step, which are encoded in a main function called “Go”. More specifically, I indicate which class of agents executes each process and which state variables are updated, strictly following the exact order of the processes as they appear in the programme. While the most basic processes are fully explained in this subsection, all details on more structured functions and information on parameters are provided in Subsection 3.3 (Submodels).

1. **Checking the number of bacteria on the culture plate.** The observer determines whether the bacteria displayed on screen should be abundant or scarce (their specific number and size are arbitrary). If bacteria are scarce, the observer updates *stored-cost-of-starvation-%*. For further details, see Subsection 3.3.
2. **Checking the presence of *C. elegans* larvae on the culture plate.** The observer determines whether *C. elegans* larvae should be present in the simulation world or not (their specific number and size are arbitrary). For further details, see Subsection 3.3.
3. **Bacterial movement and replication.** Bacteria make minimal movements and update *heading*, *xcor*, and *ycor*. Every 10 ticks (= 1 hour, which is a realistic timing for *E. coli*), each bacterium produces a new, identical bacterium by binary fission.
4. **Movement and replication of *C. elegans* larvae.** *C. elegans* nematodes execute the submodel “move-worms” and update their *heading*, *xcor*, and *ycor*. Every 10 ticks (= 1 hour, which is an arbitrary value), they produce a new *C. elegans* nematode with *size* = 0.45.
5. **Control of environmental (dietary) fluctuations.** The role of prevalent food source on the culture plate can be periodically alternated between bacteria and *C. elegans* larvae. For further details, see Subsection 3.3.
6. **Random mortality of *P. pacificus* eggs.** The observer keeps the total number of newly generated *P. pacificus* nematodes (i.e., with *developmental-stage* = “none”) below the value of *Carrying-Capacity* (see Subsection 3.3) by eliminating all exceeding new nematodes.
7. **Start of the development of *P. pacificus* eggs.** The development of the surviving eggs begins. Newly generated *P. pacificus* nematodes that survived the previous time step (the current value of *developmental-progression* is 1) update their *developmental-stage* to “new-egg” and set the value of *parent-1* = *who* (their ID number).
8. **Life history of *P. pacificus* nematodes.** *P. pacificus* nematodes move, grow, and go through several developmental stages, so they constantly update their value of *heading*, *xcor*, *ycor*, *developmental-progression* and, if they have reached some specific values of the latter state variable, they also update *developmental-stage*. Until *developmental-progression* = 240, they execute the submodel “move-eggs”, then they execute the submodel “move-worms”. When they pass from a value of *developmental-stage* to the next, they can update their *shape*, *size*, *color*, and *phenotype,* and when they are adult, they also update their *time-to-oviposition*. Adult *P. pacificus* can also update their *partner* and *parent-2*. At the end of their life cycle, hermaphrodites execute the submodel “reproduce”, through which they generate their offspring by respecting the segregation ratios of classical genetics (in the context of a population which performs both self-fertilisation and occasional cross-fertilisation). Then, they apply any reproductive costs by eliminating a specific number of their offspring. All *P. pacificus* nematodes of the previous generation die immediately after ovipositioning and applying reproductive costs to their offspring. There is no specific chronological order in the execution of single reproductive events, assignments of costs, and deaths of parents within a time step, as these processes are completely independent. However, the offspring must be produced by all parents before applying any costs, and such costs must be applied by all parents before all parents die. *P. pacificus* worms with a predatory phenotype (*phenotype* = 1) can also eat *C. elegans* larvae when they are close to one of them and update *nematodes-killed*. Details on this block of computer code and all submodels called here are provided in Subsection 3.3.
9. **Model update.** The tick counter (i.e., the counter of time steps) is updated by 1 unit and the outputs of the simulation are displayed in the NetLogo interface.

Most of the processes above were included into the model to represent the whole life cycle of multiple, fully synchronised generations of *P. pacificus*, taking into account all the fundamental aspects concerning the development, behaviour, reproduction, and genetics of this species (see Supplementary File 2). Conversely, the processes executed by food sources of *P. pacificus*, i.e., bacteria and *C. elegans* larvae, are very basic (constant replication, movement on the grid) because the only role of these agents is to provide alternative nutrients for the target organism of the study. Indeed, their presence and interplay with *P. pacificus* are relevant only insofar as they affect costs and benefits in terms of offspring produced and trigger alternative plastic responses in worms with a plastic genotype. Finally, the activities carried out by the observer are of paramount importance, as they allow for the management of temporal fluctuations in the environment (in terms of the prevailing food source) and prevent exponential growth of other classes of agents. This enables the execution of targeted and well-controlled virtual experiments.

The order of execution of the scheduled activities allows to first define the food conditions that characterise the environment at each time step. This establishes the context for the complete, synchronised, and sequential unfolding of the entire life cycle of *P. pacificus*, which respects a chronological proportion with the actual developmental steps of this animal, from the egg to reproduction. In particular, the production of a new generation of nematodes by all *P. pacificus* worms within a single time step does not follow any order of execution among agents, as each reproductive event is totally independent of all others occurring simultaneously. Nevertheless, it is important that reproduction always takes place before the application of costs in terms of offspring, and that the application of these costs always takes place before all parents die, since all these actions are executed within a single time step.

### 2. Design concepts

This model provides a simulation platform to investigate the conditions affecting the evolution of phenotypic plasticity and/or the spread of plastic/non-plastic strategies in populations, which are important evolutionary topics frequently addressed using both mathematical (Gabriel & Lynch, 1992; King & Hadfield, 2019; Kasada & Yoshida, 2020) and computational modelling (Lalejini & Ofria, 2016; Miras, 2024). At least one of the main variables implemented here (i.e., inherent cost of plasticity, timescale of environmental oscillation, and degree of plasticity) is often included in the design of works such as those cited above, since these variables are known to play a significant role in the evolution of plasticity. The bottom-up approach typical of agent-based modelling has also proven to be effective in this field, e.g., using NetLogo (Edelaar et al., 2017) or providing an integrative framework between theory and experimental data obtained from a very prominent model organism, i.e., *P. pacificus* (Kalirad & Sommer, 2024). Similarly to the latter models, *PhePlastiComp* was therefore designed to heavily rely on the principles of agent-based modelling and complex adaptive systems (CAS) (Grimm, 2005), including emergence, adaptation, objectives, prediction, sensing, interaction, and stochasticity, which are explicitly mentioned in the next paragraphs.

1. **Emergence.** Changes in the gene pool of populations of *P. pacificus*, which ultimately lead to fixation/extinction of one of the two genetic variants of *eud-1*, are emerging patterns (i.e., spontaneously occurring global behaviours of the system) generated by a multitude of individual reproductive events taking place at the end of each generation. These changes in allele frequency are not imposed by the program and can vary unpredictably, as they depend on many local factors including reproductive benefits and costs associated with the variable *genotype* or with the development of a specific *phenotype*, the level of individual responsiveness to the environment, and the timescale of food sources variation. In contrast, the fixed population size within a simulation can be viewed as an imposed outcome (due to the parameter *Carrying-Capacity*, see Subsection 3.3). This is a reasonable simplification, since monitoring variations in population size was not a focus of the model.
2. **Adaptation.** There is evidence that both mouth phenotypes of *P. pacificus* have an adaptive value. Indeed, it was demonstrated that different fitness advantages are conferred on each mouth-form depending on the alternative environmental conditions experienced by nematodes in terms of food availability, e.g., scarcity of bacteria and presence of *C. elegans* prey, and vice-versa (Serobyan et al., 2014; Susoy & Sommer, 2016). This important element of biological realism was introduced into the model by increasing the probability of each *P. pacificus* nematode to develop the most advantaged *phenotype* depending on the dietary conditions experienced during its lifetime. In other words, the irreversible developmental decision of individual *P. pacificus* agents can be considered an adaptive behaviour.
3. **Objectives.** The measure of individual success is calculated in terms of fitness, i.e., the number of offspring produced by a *P. pacificus* nematode at the end of its life cycle. Fitness was implemented into the model by assigning different benefits and costs to individual worms, depending on the value of the state variables *genotype* and *phenotype* and the prevalent food source. The number of larvae killed by worms with a predatory mouth-form (*nematodes-killed*) also plays a role in this calculation. These benefits and costs ultimately translate into percentage changes in the number of offspring produced. Thus, individuals with a plastic *genotype* actively pursue increases in their fitness by increasing the probability of developing a certain phenotype depending on the environment and, for facultative predators, by eating larvae which are closest to them on the NetLogo lattice.

4. **Prediction.** *P. pacificus* agents can predict future environmental conditions (within the same generation) based on the conditions experienced during their earlier developmental steps. The probability to develop a given *phenotype* once adulthood is reached is selected by each *P. pacificus* worm with a plastic *genotype* by foreseeing that the same conditions will persist at least until the end of the current generation. This is a precise modelling choice based on biological evidence indicating that irreversible developmental decisions are more likely to evolve when environmental conditions vary between generations, rather than within the same generation (Kassen, 2002).
5. **Sensing.** *P. pacificus* nematodes can sense directly the abundance of bacteria on the whole plate by counting the members of this agent class. The number of bacteria is arbitrary, as its only function is to distinguish a context of abundance from one of scarcity. The perceived quantity of bacterial food is crucial, because microbial abundance affects the developmental decision of *P. pacificus* agents analogously to their biological counterparts (Bento et al., 2010; Susoy & Sommer, 2016). *P. pacificus* worms can also sense the location (*xcor*, *ycor*) of close *C. elegans* larvae, as they can try to eat such prey when the distance between predator and prey is smaller than the size of a single NetLogo patch. To form pairs, male *P. pacificus* worms can also sense the value of the state variables *sex* and *partner* of all conspecifics on the plate. If a male finds a random hermaphrodite whose literal value of the variable *partner* is “nobody”, he can impose that hermaphrodite to set himself (*who* number) as *partner*. Finally, newly generated *P. pacificus* agents have access to all state variables of their parent (also including the variable *parent-2* of their father, if they are the result of a cross), so they can decide which characteristics to inherit (e.g., *species* or, in accordance with the rules of genetic transmission, *genotype*) and which to modify (e.g., *size*, *shape*, and *phenotype*).
6. **Interaction.** Interaction between *P. pacificus* and its food sources is a fundamental feature of the model, as it affects both developmental decisions (*phenotype*) and individual fitness. The nutritional/predatory interaction is mediated in the case of bacteria, whereas it is direct, based on physical proximity between agents, in the case of *C. elegans* larvae. Mediated interaction was also used to model pair formation, which results in hermaphrodite x male crosses, since there is no mate choice and thus there was no need to implement direct contact between partners. In contrast, direct interaction is critical for ovipositioning, since each hermaphrodite nematode is responsible for producing and laying its eggs in the exact position in which it is located at the end of its life cycle, which is also an element of biological realism. Additional information on all these interactions is provided in the previous paragraph dedicated to sensing.
7. **Stochasticity.** Stochastic processes included in the model rely on NetLogo primitives based on the Mersenne Twister algorithm, which generates uniform pseudo-random numbers with a very long period (Matsumoto & Nishimura, 1998). While such processes are a clear simplification in the case of events that are not central to the investigation carried out here (e.g., the maintenance of population size below the selected *Carrying-Capacity* through the application of random mortality to *P. pacificus* eggs), in other cases it is an element that is consistent with current empirical data. For instance, stochasticity is involved in events of meiotic non-disjunction leading to the production of males (Hong & Sommer, 2006), and both a stochastic and a conditional component are involved in the developmental decision leading to the formation of the definitive *phenotype* in adult nematodes (Susoy & Sommer, 2016).
8. **Observation.** The evolution of digital populations of *P. pacificus* is monitored by the NetLogo simulation platform by reporting, at each time step, the current *generation*, the number of *P. pacificus* nematodes for each adult *phenotype* (bar chart), the relative frequency of each adult *phenotype* (numerical monitors), the relative frequencies of each *genotype* and allele (numerical monitors and line plots), and the relative frequency of plastic vs non-plastic genotypes (numerical monitors). In this study, I focus on the values of the state variables *genotype* and *generation* at the final time step of each run, which I used to obtain data on the frequency of fixation and time until fixation of the two alleles involved (i.e., wild-type and mutant *eud-1*). For times until fixation, I used the outputs of runs to obtain both measures of central tendency (mean and median) and measures of dispersion of data (range, interquartile range). The final time step of each simulation corresponds to the first tick where only one of the two alleles is still present in the gene pool following the extinction of the other allele.

### 3. Details

#### 3.1 Initialisation

All the instructions necessary for initialisation, i.e., to create all entities and set the initial values of their state variables, were implemented through a NetLogo procedure called “Setup”. Here I describe in detail the algorithm contained within this procedure (see Subsection 3.3 for a description of all parameters mentioned in the list below. To distinguish them from state variables, I used capital letters for their initials).

1. The simulation world is cleared of all elements from the previous simulation (if any).
2. All patches turn light blue (*pcolor* = 89).
3. The state variable *stored-cost-of-starvation-%* takes on the value of the parameter *Cost-of-Starvation-%*. This allows the model to store the value of the individual cost of starvation, which is applied when microbial food is scarce, so the same cost can be reapplied in future phases of microbial starvation.
4. If *Abundant-Bacteria?* = TRUE, 800 bacteria with *shape* = “circle”, *color* = 46, *size* = 0.45, and a random value of *xcor* and *ycor* are created. If *Abundant-Bacteria?* = FALSE, 200 bacteria with identical characteristics as above are created.
5. If *C.elegans?* = TRUE, 300 *C. elegans* nematodes with the *shape* of a worm (custom shape), *color* = grey, *size* = 0.45, a random value of *xcor* and *ycor*, and *species* = “c.elegans” are created. If *C.elegans?* = FALSE, 0 *C. elegans* nematodes are created.
6. A specific number of wild-type homozygous (*++*) *P. pacificus* nematodes is created based on the value assigned to the parameter *Initial-++*. These nematodes have *shape* = “circle”, *color* = grey, *size* = 0.5, *species* = “p.pacificus”, *genotype* = “++”, *phenotype* = 0, *sex* = “hermaphrodite”, *partner* = nobody, *developmental-stage* = “none”, *developmental-progression* = 0, *generation* = 0, *parent-1* = *who* (ID number), *parent-2* = nobody, and *nematodes-killed* = 0.
7. A specific number of heterozygous (*+e*) *P. pacificus* nematodes is created based on the value assigned to the parameter *Initial-+e*. These nematodes have the same characteristics as those described in point 6, except for their *genotype* (= “+e”).
8. A specific number of mutant homozygous (*ee*) *P. pacificus* nematodes is created based on the value assigned to the parameter *Initial-ee*. These nematodes have the same characteristics as those described in the previous two points, except for their *genotype* (= “ee”).
9. A specific number of wild-type hemizygous (*+O*) *P. pacificus* nematodes is created based on the value assigned to the parameter *Initial-+O*. These nematodes have the same characteristics as those described in the previous three points, except for their *genotype* (= “+O”) and their *sex* (= “male”).
10. A specific number of mutant hemizygous (*eO*) *P. pacificus* nematodes is created based on the value assigned to the parameter *Initial-eO*. These nematodes have the same characteristics as those described in point 9, except for their *genotype* (= “eO”).
11. The time step (tick) counter is set/reset to 0.

The model aims to explore the evolution of the simulated system across several scenarios which differ from each other only in terms of the initial conditions used. Since I was interested in investigating the effects of different genetic compositions of the initial population, the only parameters used in this step (other than those regulating the presence/absence of *E. coli* and *C. elegans*, *Abundant-Bacteria?* and *C.elegans?*, and the cost associated with starvation, *Cost-of-Starvation-%*) are those concerning the initial number of *P. pacificus* nematodes (eggs) for each genotype. Consequently, the state variable *genotype* is particularly important, and the initial genotype ratio of the population must change depending on the allele frequency considered for each of the two variants of *eud-1*, i.e., *+* (wild-type) and *e* (mutant).

#### 3.2 Input data

One of the crucial features of the model is the presence of periodic environmental fluctuations, expressed in terms of reversions in the prevailing food source (*E. coli* or *C. elegans*) during simulations. This function was implemented using two parameters named *Fluctuating-Environment?* and *Environmental-Cycles*, which affect the two switch-like environmental variables *Abundant-Bacteria?* and *C.elegans?* by turning them on (TRUE) and off (FALSE) alternately, starting from the value assigned to each of them during initialisation. More specifically, *Fluctuating-Environment?* is another Boolean variable which, when set to TRUE, makes the environment fluctuate over simulation time, whereas *Environmental-Cycles* can take on any positive integer value and represents the time interval, expressed in number of non-overlapping generations of *P. pacificus*, at the end of which the two variables *Abundant-Bacteria?* and *C.elegans?* reverse their value. In this study, simulations were always initialised with opposite values of the two variables indicating the prevalent nutrient source, in order to represent the alternation of dietary regimes.

Although this approach to managing environmental variability is of course a simplification, as it assumes that relevant changes always occur at regular intervals, it may also reflect the real trends of a wide range of biological systems, which are often characterised by periodic environmental oscillations at the most disparate scales (Abdul-Rahman et al., 2021). Another limitation is the impossibility of simulating within-generation changes, but since the model focuses on an organism whose life cycle includes an irreversible developmental decision, it is reasonable to assume that simulating between generations changes is appropriate for this case study (Kassen, 2002). As for the timescales explored in my scenarios, i.e., with *Environmental-Cycles* = 1, 5, 10, 15, and 20, they were chosen because they could match data obtained from real ecosystems where *P. pacificus* can be found. For instance, in carcasses of beetles co-infested by other nematodes, mouth-form ratios can undergo major environmentally-induced changes within a few weeks, that is, across several (but not many) generations of worms (Renahan & Sommer, 2022). Keeping in mind the heterogeneity of *P. pacificus* habitats, which include soil, scarab bodies and corpses, and decomposing vegetal matter (Susoy & Sommer, 2016; Félix et al., 2018), it is also plausible that, in some cases, a specific nutrient source may be more prevalent than another (on average) for a time interval spanning several generations (although it is also likely that nematodes of this species may experience unpredictable and/or faster environmental change in other circumstances).

#### 3.3 Submodels

This subsection is primarily dedicated to providing a full description of all functions (i.e., NetLogo procedures) executed by agents at each single iteration, as they are called within the main function/procedure of the program (“Go”). I also describe in greater detail the parts of the main procedure which have not been fully examined in Subsection 1.3 (Process overview and scheduling), particularly those concerning the life history of *P. pacificus* nematodes. However, to gain a proper understanding of this description, I first present all the parameters used during simulations and provide an explanation/rationale for each of them. The outcomes of preliminary tests and simulation scenarios performed to calibrate some of the parameters and validate the global behaviour of the model are presented within the Results and Discussion section of the main text.

##### 3.3.1 Parameters

All parameters are summarised, described, and justified in Table S2. While most parameters are discussed in the following paragraphs, the parameters named *Abundant-Bacteria?*, *C.elegans?*, *Fluctuating-Environment?*, and *Environmental-Cycles* have already been discussed in Subsection 3.2 (Input data).

**Table S2. Summary of parameters.** Parameters implemented in *PhePlastiComp* and used during simulations. This list includes the type of parameter, the range of values used in the simulated scenarios, a short description, and references supporting the implementation of each parameter and the considered values (if relevant).

| **Parameter** | **Type** | **Range** | **Description** | **References** |
| --- | --- | --- | --- | --- |
| *Abundant-Bacteria?* | Boolean | FALSE/TRUE | Quantity of bacteria on the culture plate (low/high quantity) | (Serobyan et al., 2014) |
| *C.elegans?* | Boolean | FALSE/TRUE | Presence of *C. elegans* larvae on the culture plate (not present/present) | (Serobyan et al., 2014) |
| *Fluctuating-*  *Environment?* | Boolean | FALSE/TRUE | It indicates whether dietary conditions are fluctuating over time (fixed/fluctuating environment) | (Abdul-Rahman et al., 2021; Lalejini et al., 2021) |
| *Environmental-*  *Cycles* | Integer number | 1-20 | Number of *P. pacificus* non-overlapping generations at the end of which the proportions of the two alternative food sources are reversed (i.e., the values of the parameters *Abundant-Bacteria?* and *C.elegans?* are reversed) | (Abdul-Rahman et al., 2021; Renahan & Sommer, 2022) |
| *Carrying-Capacity* | Integer number | 100-300 | Maximum population size allowed (i.e., random extrinsic mortality removes exceeding *P. pacificus* nematodes immediately after being generated by their parents) | - |
| *Probability-of-Non-*  *Disjunction-%* | Integer number | 1 | Probability to generate a male *P. pacificus* offspring due to events of meiotic non-disjunction | (Hong & Sommer, 2006; Schroeder, 2021) |
| *Degree-of-*  *Plasticity-%* | Integer number | 0-20 | Increase/decrease in the individual probability of developing the mouth phenotype that is adaptive under the experienced dietary conditions (the baseline probability is close to that observed in strain PS312 of *P. pacificus* under standard laboratory conditions). This is only applied to plastic genotypes of *P. pacificus* (*++*, *+e*, and *+O*) | (Susoy & Sommer, 2016; Crowther et al., 2023) |
| *Cost-of-*  *Plasticity-%** | Integer number | 0-3 | Fitness cost in terms of offspring produced by a parent (or a couple of parents) with a plastic genotype (*++*, *+e*, and *+O*). This is related to the individual ability to express a plastic response to dietary conditions | (DeWitt et al., 1998; Murren et al., 2015; Dardiry et al., 2023) |
| *Cost-of-*  *Starvation-%** | Integer number | 0-3 | Fitness cost in terms of offspring produced by a parent (or a couple of parents) during time intervals when bacteria are scarce | (Serobyan et al., 2014; Werner et al., 2017; Mautz et al., 2019) |
| *Benefit-of-Faster-*  *Development-%** | Integer number | 0-3 | Fitness benefit in terms of offspring produced by a parent (or a couple of parents) with a ST phenotype (*phenotype* = 2), if bacteria are abundant | (Serobyan et al., 2014; Sommer et al., 2017; Dasgupta et al., 2022) |
| *Benefit-of-*  *Predation-%** | Integer number | 0-3 | Fitness benefit in terms of offspring produced by a parent (or a couple of parents) with an EU phenotype (*phenotype* = 1). The outcome is proportional to the number of prey eaten during adulthood | (Serobyan et al., 2014; Sommer et al., 2017) |
| *Efficiency-of-*  *Attack-%* | Integer number | 8 | Probability to kill a close prey. The number of killed prey (up to a maximum of 5) must be multiplied by *Benefit-of-Predation-%* to obtain the total benefit for that specific nematode | (Serobyan et al., 2014) |

*The baseline number of offspring, i.e., the offspring produced by each hermaphrodite *P. pacificus* nematode without considering any costs or benefits, is 100.

***Carrying-Capacity.*** It is used to keep the population size below a certain value by applying a random extrinsic mortality to the offspring of all *P. pacificus* nematodes immediately after egg laying. In this study, I kept this value fixed at 300, which is a reasonable limit so as not to provoke overcrowding within the virtual culture plate.

***Probability-of-Non-Disjunction-%.*** *P. pacificus* nematodes are either hermaphrodites (XX) or hemizygous males (XO). Males may be introduced from the start of simulations, but they may also be generated during simulations due to meiotic non-disjunction, with a given probability per reproductive event which is regulated by the parameter *Probability-of-Non-Disjunction-%*. This spontaneous arise of new males is very infrequent both in reality, where males are usually less than 1% of the population (Hong & Sommer, 2006; Schroeder, 2021), and in my simulations. Almost all reproductive events occurring in the experimental scenarios consist of self-fertilisation/selfing (because most nematodes are hermaphrodites), with only a small fraction of crosses between hermaphrodites and hemizygous males.

***Degree-of-Plasticity-%.*** The reason behind the inclusion of this variable is that discrete plastic responses (polyphenisms), including the developmental decision of *P. pacificus*, may show a different level of plasticity depending on the steepness of their switch-like reaction norm (Crowther et al., 2023). This parameter enables the regulation of the individual probability of developing an adaptive mouth-form under each dietary condition. This means that adjusting the value of *Degree-of-Plasticity-%* implies either an increase or a reduction in such probability for all plastic genotypes (*++*, *+e*, and *+O*). When *Degree-of-Plasticity-%* = 0, the probability that each plastic genotype develops a given mouth-form reflects the values reported under standard laboratory conditions (which include abundant bacterial food and no overcrowding (Susoy & Sommer, 2016)) for wild-type and mutagenised genotypes of the reference strain PS312 (Ragsdale et al., 2013). Any increase in the value of this parameter moves plastic genotypes away from these baseline, characteristic values of the reference strain, making such genotypes more plastic. This makes the model suitable for the exploration of different levels of environmental responsiveness.

***Cost-of-Plasticity-%.*** This cost in terms of individual fitness is only applied to plastic genotypes (*++*, *+e*, and *+O*) and refers to the number of random zygotes/newly generated eggs eliminated from the grid for each parent (or couple of parents) belonging to a plastic strain. Here, the cost is associated with the ability to respond to environmental cues like diet or starvation by increasing the probability of developing a particular phenotype and/or, more generally, with the capacity to make a developmental decision and retain the dimorphism. This inherent cost of plasticity may be related to the production of the plastic response or to the maintenance of sensory or regulatory elements which are not required by non-plastic genotypes (DeWitt et al., 1998; Murren et al., 2015; Dardiry et al., 2023). In crosses, *Cost-of-Plasticity-%* is applied two times if both partners belong to a plastic strain.

***Cost-of-Starvation-%.*** It is a reduction in fecundity applied to all *P. pacificus* nematodes if microbial food is scarce and refers to the number of random zygotes/newly generated eggs removed from the grid for each parent (or couple of parents) experiencing starvation. As a matter of fact, bacteria are by far the best source of nutrients for both morphs (Serobyan et al., 2014) and it was ascertained that the complete absence of bacteria leads to slower development and delayed sexual maturation (Werner et al., 2017). Although dietary restriction is not directly harmful for parents in other nematode species (under certain conditions), it was found that it can rather reduce the fitness of offspring (Mautz et al., 2019). Thus, for simplicity, in the model this reduction in fitness is directly applied to parents experiencing bacterial starvation. As for crosses, this cost is applied two times, regardless of the genotype of the two partners.

***Benefit-of-Faster-Development-%.*** It is an increase in the number of offspring applied each time to a different, random genotype among the zygotes produced by a ST nematode when bacteria are abundant. For example, if *Benefit-of-Faster-Development-%* is set to 2, microbial food is abundant, and we consider a ST nematode with *+e* genotype performing selfing, the additional number of offspring (2) will be applied either to *++*, *+e*, or *ee* zygotes from case to case. In crosses, *Benefit-of-Faster-Development-%* is applied only one time, and only if both partners are ST. In fact, since the fitness advantage depends on the anticipation of sexual receptivity, this anticipation needs to be synchronised in both parents for the advantage to be exploited. The increase in fitness for ST worms due to their faster developmental rate is reported in literature (Serobyan et al., 2014). In the model, it was implemented as an increase in fecundity and not as a difference in life cycles to keep all developmental steps synchronised for both forms. The application of this parameter is also supported by comparisons with similar dynamics in other ecdysozoans, such as *Drosophila melanogaster*, where the anticipation of sexual maturation under certain dietary conditions may correspond to an early-life increase in the number of offspring (Dasgupta et al., 2022).

***Benefit-of-Predation-%.*** It is an increase in the number of offspring applied each time to a different, random genotype among the zygotes produced by an EU nematode when feeding on *C. elegans* larvae. This value must be multiplied by the sum of larvae killed by the EU nematode during its life cycle (up to a maximum of 5 larvae, so as not to confer unrealistic advantages). The number of prey killed depends on the additional parameter ***Efficiency-of-Attack-%***, that is the probability to kill a close larva. For instance, if *Benefit-of-Predation-%* is set to 2 and an EU hermaphrodite with *+e* genotype kills 2 prey, this additional number of offspring (i.e., 2*2 = 4) will be applied either to *++*, *+e*, or *ee* zygotes from case to case. In crosses with other EU nematodes, *Benefit-of-Predation-%* must be multiplied by the sum of all larvae killed by both partners during their life cycles. This increase in fecundity was also observed experimentally. Indeed, EU worms clearly showed their superiority in killing prey over the other form, supporting the hypothesis of an advantage in fitness on a mixed diet (Serobyan et al., 2014).

##### 3.3.2 Checking the number of bacteria submodel

Here I describe the submodel that regulates the quantity of bacteria on the culture plate. This is a block of computer code contained within the main procedure (“Go”).

If *Abundant-Bacteria?* = TRUE, the observer keeps the number of bacteria strictly below 800 and sets the parameter *Cost-of-Starvation-%* = 0. If *Abundant-Bacteria*? = FALSE, the observer keeps the number of bacteria strictly below 200 and sets *Cost-of-Starvation-%* = *stored-cost-of-starvation-%*. The value of the latter state variable does not change during a simulation.

##### 3.3.3 Checking the presence of *C. elegans* larvae submodel

Here I describe the submodel that regulates the presence of *C. elegans* larvae on the culture plate. This is a block of computer code contained within the main procedure (“Go”).

If *C.elegans?* = TRUE, if some *C. elegans* nematodes are already present, the observer keeps the number of *C. elegans* nematodes strictly below 300, otherwise the observer creates 300 new *C. elegans* nematodes. If *C.elegans?* = FALSE, all *C. elegans* nematodes die.

##### 3.3.4 Control of dietary fluctuations submodel

Here I describe the submodel that regulates the alternation of dietary regimes on the culture plate. This is a block of computer code contained within the main procedure (“Go”).

If the parameter *Fluctuating-Environment?* = TRUE, the current time step is not 0, and the current time step is a multiple of the product between the duration of a generation of *P. pacificus* (i.e., 820 ticks = 82 hours) and the value of *Environmental-Cycles*, the observer reverses the binary value of *Abundant-Bacteria?* and that of *C.elegans?*. Consequently, the prevalent food source is periodically switched between bacteria and *C. elegans* larvae.

##### 3.3.5 Life history submodel

Here I describe in detail the submodel that enables *P. pacificus* agents to complete their entire life cycle. Similarly to the subsections above, this block of computer code is not a submodel in the strict sense, but rather a very substantial sequence of instructions located in the main procedure (the “Go” procedure). It is also described in general terms in Subsection 1.3 (Process overview and scheduling) and contains three proper submodels, which are described in the following subsections: “move-eggs”, “move-worms”, and “reproduce”.

In this portion of the programme, the observer asks all *P. pacificus* nematodes to update *developmental-progression* by 1 unit. If *developmental-progression* = 150, nematodes set *developmental-stage* = “J1” and set *size* = 0.75. If *developmental-progression* < 240, nematodes execute the submodel “move-eggs”, otherwise they execute the submodel “move-worms”. If *developmental-stage* = “J1” and *developmental-progression* = 200, nematodes set *developmental stage* = “J2” and *size* = 1. If *developmental-stage* = “J2” and *developmental-progression* = 240, they set *shape* “nematode” (custom shape) and *size* = 4. If *developmental-progression* = 400, they set *developmental-stage* = “J3” and size = 5. If *developmental-progression* = 480, they set *developmental-stage* = “J4” and *size* = 6.

If *developmental-progression* = 700, they set *developmental-stage* = “adult” and, depending on their *genotype* and the value generated by a function that calculates the probability of developing the mouth-form EU (see Subsection 3.3.9, Custom reporters), they decide either to set *phenotype* = 1 (EU), the custom *shape* “EU”, and *color* blue, or *phenotype* = 2 (ST), the custom *shape* “ST”, and *color* red. For instance, if their *genotype* is “++”, they have a genotype-dependent probability to develop into EU worms, otherwise they become ST worms. Of course, this is only for plastic genotypes (“++”, “+e”, and “+O”). Non-plastic genotypes (“ee” and “eO”) always set their *phenotype* = 2 (ST), *shape* “ST”, and *color* red. At this point, all *P. pacificus* nematodes set *time-to-oviposition* = 120 and *size* = 7, and if their *sex* is “male”, they search for an available hermaphrodite partner: if there is at least one “hermaphrodite” nematode with *partner* = nobody, they set one of those nematodes as their *partner* and ask their partner to set themselves (i.e., males) as their *partner* and *who* of their partner as *parent-2*.

If *developmental-progression* is strictly greater than 700 but less than or equal to 820, all *P. pacificus* nematodes decrease their value of *time-to-oviposition* by 1 unit. If their *sex* is “hermaphrodite” and *time-to-oviposition* = 0, they execute the submodel “reproduce”. After that, nematodes with *sex* = “hermaphrodite” and *developmental-progression* = 820 apply costs in fecundity (i.e., *Cost-of-Starvation-%* and/or *Cost-of-Plasticity-%*) to their own offspring, depending on the *genotype* of the only parent (if its *partner* is nobody) or on both genotypes of parents (if its *partner* is not nobody). After applying these exact costs to their offspring, all these hermaphrodite nematodes immediately die. Here is an example of the application of this cost in terms of offspring to *+e* hermaphrodites performing self-fertilisation: they ask a precise number of other nematodes in the same patch with *developmental-stage* = “none” and *parent-1* = their value of *parent-1*, where this number of other nematodes corresponds to the sum of *Cost-of-Plasticity-%* and *Cost-of-Starvation-%*, to die. In contrast, *++* hermaphrodites crossing with *+O* males will execute the following instructions: they ask a precise number of other nematodes in the same patch with *developmental-stage* = “none”, *parent-1* = their value of *parent-1*, and *parent-2* = their value of *parent-2*, where this number of other nematodes corresponds to 2 * *Cost-of-Plasticity-%* + 2 * *Cost-of-Starvation-%*, to die. Both these costs are paid two times because they also affect the fitness of the male partner (other than the hermaphrodite parent). Further details on the application of these costs are provided in Subsection 3.3.1 (Parameters). Male nematodes with *partner* = nobody and *developmental-progression* greater or equal to 820 die.

If *developmental-progression* ≥ 700, *phenotype* = 1 (EU) and *nematodes-killed* < 5, if there is at least one nematode of the species “c.elegans” within range of a patch, there is a probability corresponding to the value of *Efficiency-of-Attack-%* to ask one of *C. elegans* nematodes within range of a patch to die, which causes an increase of the value of *nematodes-killed* by 1 unit.

**Rationale and clarifications.** Worms of each generation start as eggs, then go through all developmental phases reported for this species, with a timing approximately corresponding to that observed empirically (Hong & Sommer, 2006). More specifically, all *P. pacificus* nematodes reach the J1 (i.e., Juvenile 1) stage 150 ticks (i.e., 15 hours) after egg formation, then undergo the first moult while they are still inside the eggshell, 200 ticks after egg formation, reaching the J2 stage. Hatching takes place 240 ticks after egg formation. The following steps are the J3 and J4 stages, 400 ticks and 480 ticks after egg formation, respectively. At the end of the J4 phase, after the 4^th^ moult (700 ticks after egg formation), the adult age is reached and all nematodes develop their mouth form irreversibly (Bento et al., 2010; Ragsdale et al., 2013), which allows them to feed on bacteria and/or *C. elegans* larvae. Adult worms can also reproduce, either by selfing or by crossing. In the model, each male (if present) can copulate with a single, randomly chosen hermaphrodite, while all other hermaphrodites perform selfing. 820 ticks after egg formation, i.e., 82 hours, that corresponds to the real lifespan of *P. pacificus* (Hong & Sommer, 2006; Sommer et al., 2017), all hermaphrodites produce their eggs.

Mouth dimorphism and the related behavioural plasticity in dietary habits are critical characteristics of *P. pacificus* nematodes. In a few words, EU nematodes (blue colour in the model) have large mouths and can feed both on bacteria and other nematodes, whereas ST nematodes (red colour in the model) have narrow mouths and can only feed on bacteria. Some nematodes have the potential to express either form (i.e., they have a plastic genotype), while others express only one form constitutively (i.e., they have a non-plastic genotype). In real worms, one of the main regulators of the plastic response is the sulfatase-encoding *eud-1* gene, a dosage-dependent developmental switch that is sensitive to several environmental factors and is located on the X chromosome (Ragsdale et al., 2013; Sommer et al., 2017; Werner et al., 2017; Sieriebriennikov et al., 2018; Dardiry et al., 2023). While wild-type individuals of the reference strain PS312 (*++*) express both mouth forms (on average, 80% EU and 20% ST under standard laboratory conditions with no food deprivation or overcrowding), mutations affecting this key gene cause changes in phenotype ratios (Susoy & Sommer, 2016). This was observed with loss-of-function mutations producing dominant *eud-1* (in the model, *e*) alleles, that induce a shift towards ST-biased outputs to the point of making mutants completely deficient in developing the EU form (Sommer et al., 2017; Casasa et al., 2023).

In particular, in the equivalent strain of PS312 named RS2333 (Serobyan et al., 2013; Lenuzzi et al., 2021), *ee* homozygotes (in the model, also *eO* males) lose their ability to express plastic responses, since they can only develop the ST morph, whereas *+e* heterozygotes show incomplete dominance and are still plastic, as well as *+O* males. *+e* and *+O* worms are roughly 10-40% EU in the lab (Ragsdale et al., 2013; Sommer et al., 2017). All phenotype ratios shown by *P. pacificus* agents of *PhePlastiComp* under the standard protocol are close to those observed in the lab under the same conditions. The simultaneous expression of both morphs in isogenic populations carrying at least one copy of *+* and living in a stable environment is an example of stochastic regulation of plasticity, that is typical of this species (Susoy & Sommer, 2016; Sieriebriennikov & Sommer, 2018). Both in the model and in the real world, mutants lacking the *+* allele are unable to express the dimorphism. In this regard, it is important to note that the *eud-1* mutants of the model are produced by any hypothetical mutation affecting the *eud-1* gene and having exclusive effects in the reduced (*+e*, *+O*) or null (*ee*, *eO*) expression of the EU phenotype.

In addition to genetic makeup and stochasticity, mouth-form ratios are also affected by changes in environmental conditions. Indeed, mouth dimorphism is regulated by environmental factors, the most relevant of which are starvation, population density (Susoy & Sommer, 2016), and temperature. For instance, temperatures below 15°C reduce the percentage of worms developing the EU morph in PS312 (Lenuzzi et al., 2021), and crowding is positively related to a higher expression of the EU phenotype due to a specific pheromone signal (Bento et al., 2010). However, my study considers neither the influence of temperature, as I assume to keep it fixed over time at 20°C, nor the effects of population density in mouth formation. In fact, it has been found that the pheromones inducing shifts towards more EU-biased ratios in juvenile individuals are stage specific and come from adults (Werner et al., 2018). Since generations are rigorously non-overlapping in the model, juvenile *P. pacificus* worms are never in contact with adults, so adults’ pheromones cannot influence the irreversible developmental decision of younger nematodes. Consequently, *PhePlastiComp* was designed to apply environmentally-induced shifts in phenotype ratios only in response to dietary variation.

A reduced presence of bacteria in the simulation grid induces an increase (*Degree-of-Plasticity-%*) in the probability of developing the EU form for each plastic genotype, whereas non-plastic genotypes are not responsive when subject to changes in food availability. In other words, the parameter *Degree-of-Plasticity-%* represents the level of conditional plasticity, and it generates variation in the EU/ST ratio in response to changes in food availability (for further details on the level of plasticity of switch-like reaction norms, see Crowther et al. (2023)). It is an important variable because it enables the investigation of different levels of environmental responsiveness, including the extreme case (0%) where regulation of plasticity is only stochastic and not conditional. The increase in EU frequency in plastic strains in response to low bacterial food is well-acknowledged (Susoy & Sommer, 2016) and represents a case of adaptive plasticity. Indeed, the ability to express the EU morph enables the exploitation of alternative food sources, such as nematodes of other species (in the model, *C. elegans* larvae), resulting in greater fitness for that environmentally-induced phenotype under such conditions of microbial scarcity (Serobyan et al., 2014).

As for the possible formation of stress resistant dauer larvae due to prolonged periods of food deprivation (Werner et al., 2018), I excluded this developmental arrest from the model by allowing the presence of reduced quantities of bacteria in all runs. Indeed, in the real world, bacteria may produce dauer-inhibitor chemical(s) that antagonise this process (Ogawa & Brown, 2015).

All information about mouth-form ratios for each genotype and under each environmental context in *PhePlastiComp* is summarised in Table S3. The correct application of these genotype- and environmental-dependent probabilities of developing each phenotype was confirmed by preliminary runs aimed at inspecting i) the relevant state variables (*genotype*, *phenotype*, *sex*, *age*) of single *P. pacificus* agents under each dietary condition, and ii) the phenotype ratio of the entire population, by monitoring the bar chart showing the frequency of mouth forms in the interface.

**Table S3.** Mouth-form ratios in relation to genotype and variation in food sources.

| **Genotype** | **Frequency of EU with abundant bacteria** | **Frequency of ST with abundant bacteria** | **Frequency of EU with scarce bacteria** | **Frequency of ST with scarce bacteria** |
| --- | --- | --- | --- | --- |
| *++*, wild-type homozygotes, plastic | ≈ 80% | ≈ 20% | ≈ 80% + *Degree-of-Plasticity-%* | ≈ 20% - *Degree-of-Plasticity-%* |
| *+e*, heterozygotes, plastic | ≈ 25% | ≈ 75% | ≈ 25% + *Degree-of-Plasticity-%* | ≈ 75% - *Degree-of-Plasticity-%* |
| *ee*, mutant homozygotes, non-plastic | 0% | 100% | 0% | 100% |
| *+O*, wild-type males, plastic | ≈ 25% | ≈ 75% | ≈ 25% + *Degree-of-Plasticity-%* | ≈ 75% - *Degree-of-Plasticity-%* |
| *eO*, mutant males, non-plastic | 0% | 100% | 0% | 100% |

##### 3.3.6 “move-eggs” submodel

*P. pacificus* eggs slowly fluctuate around the simulation world. More specifically, they move forward by one tenth of a patch and change their *heading* clockwise by an angle whose amplitude is equal to the value of a random number between 0 and 29 and anticlockwise following the same rule.

##### 3.3.7 “move-worms” submodel

*P. pacificus* worms actively move around the simulation world. In particular, they move forward by a quarter of a patch and change their *heading* clockwise by an angle whose amplitude is equal to the value of a random number between 0 and 14 and anticlockwise following the same rule.

##### 3.3.8 “reproduce” submodel

Adult *P. pacificus* nematodes at the end of their life cycle produce their offspring by following segregation ratios consistent with monohybrid inheritance and applying benefits in fitness, i.e., adding a specific number of offspring to the baseline number (100). All possible reproductive events that may occur during simulations, both in the case of self-fertilisation and crossing, are contemplated within this submodel, that is therefore made up of a series of small blocks of computer code with instructions that are largely repeated identically, except for some critical details.

The highest-level decision in this submodel is based on whether a male partner is present or not. If *partner* = nobody, nematodes must make decisions based solely on their *genotype* (“++”, “+e”, or “ee”). Then, they must evaluate whether *phenotype* = 2 (ST) and *Abundant-Bacteria?* = TRUE. If both conditions are met, a number of offspring equal to the value of *Benefit-of-Faster-Development-%* will be added to the total offspring number. Otherwise, the value resulting from *Benefit-of-Predation-%* * *nematodes-killed* will be added to the total offspring number (note that, for EU worms living in a prey-free environment and ST worms, *nematodes-killed* = 0, so the above conditions alone can represent all possible situations). The baseline number of offspring is always 100, and each benefit in terms of offspring number is applied by each hermaphrodite parent with the same probability to only one of the resulting offspring genotypes. Genotype ratios of offspring are exactly those expected based on the *genotype* of the parent, i.e., 100 *++* offspring for *++* parents, 100 *ee* offspring for *ee* parents, and 25 *++*, 50 *+e*, and 25 *ee* offspring for *+e* parents (for simplicity, no random fluctuation around these values was considered). Except for the value of *genotype* (or both *sex* and *genotype*, depending on *Probability-of-non-Disjunction-%*, which can lead to the production of *+O* or *eO* males), the characteristics of offspring are always the same: *shape* = “circle”, *color* = grey, *size* = 0.5, *phenotype* = 0, *sex* = “hermaphrodite”, *developmental-stage* = “none”, *developmental-progression* = 0, *generation* = *generation* + 1, and *nematodes-killed* = 0.

The total number of offspring (*n_o_*) resulting from self-fertilisation of each digital *P. pacificus* parent is given by Equation (1):

$$n_{o}=100-[(1-i_{ab})\cdot C_{s}]-(i_{pl}\cdot C_{P})+(i_{EU}\cdot B_{pr}\cdot n_{k})+(i_{ST}\cdot B_{fd}\cdot i_{ab})$$

(1)

, where *C_s_* is the *Cost-of-Starvation-%*, *C_P_* is the *Cost-of-Plasticity-%*, *B_pr_* is the *Benefit-of-Predation-%*, *n_k_* is the number of *C. elegans* prey killed by the parent (*nematodes-killed*), *B_fd_* is the *Benefit-of-Faster-Development-%*, and $i_{x}\in\left\{ 0, 1 \right\}$ are Boolean variables indicating environmental or phenotypic states of the parent, i.e., abundance of microbial food (*Abundant-Bacteria?*, *i_ab_*), plastic genotype (*i_pl_*), EU phenotype (*i_EU_*), and ST phenotype (*i_ST_*).

The same combinatorial logic is used to model all possible reproductive events involving a male partner (i.e., when *partner* is not nobody). In these cases, hermaphrodite nematodes must set their *nematodes-killed* counter = the sum of nematodes killed by themselves and those killed by their *partner*. The genotype ratio of offspring will be calculated based on the *genotype* of both parents, and hermaphrodite parents must evaluate whether *phenotype* = 2 (ST), *phenotype* of partner = 2, and *Abundant-Bacteria?* = TRUE to apply the correct benefit in terms of offspring produced. The characteristics of the offspring are the same as above, except that in this case, for simplicity, the possibility of further non-disjunction events is not considered. More generally, in the context of the simulations I carried out, crosses are rare events and therefore constitute an additional element of realism that could potentially be overlooked.

**Rationale and clarifications.** All selfing events and crosses, which generate 100 zygotes if no benefits are applied, were modelled as simple applications of the Mendelian patterns of inheritance. For instance, a *+e* hermaphrodite performing selfing (*+e* x *+e*) will generate 100 offspring with the following genotype ratio: ¼ *++* : ½ *+e* : ¼ *ee*, that is, 25 wild-type homozygotes, 50 heterozygotes, and 25 mutant homozygotes. A small fraction of these zygotes can also be *+O* or *eO* males, depending on the value of the parameter *Probability-of-Non-Disjunction-%* (Schroeder, 2021). Likewise, a *ee* x *+O* cross will respect the following ratio: ½ *+e* : ½ *+O*, that is, 50 heterozygotes, and 50 wild-type hemizygous males (for simplicity, non-disjunction events are not applied to crosses). However, most zygotes from each reproducing worm are eliminated after fertilisation due to random mortality, and benefits and costs in terms of fitness must be considered to obtain the real number and genotype ratio of offspring.

Mouth-form plasticity is preserved in *P. pacificus* due to context-dependent benefits in fitness for each alternative morph (Serobyan et al., 2014). Together with some costs (see Subsection 3.3.5), these benefits are introduced into the model in terms of variation in fecundity, i.e., by adding or subtracting a specific quantity from the total number of zygotes generated by each reproducing nematode. Variations in fecundity also depend on the genotype of the parent(s) and the environmental conditions experienced before reproduction. Although I explored a variety of values for these parameters in my scenarios, I always used low or moderate costs and benefits in terms of fitness (1-5%) so as not to assign unrealistically high losses or gains to agents.

The following examples should make it clearer the general functioning of the reproductive dynamics implemented in *PhePlastiComp*. For instance, consider a plate rich in bacteria and a ST hermaphrodite with a *+e* genotype performing selfing. If *Cost-of-Plasticity-%* = 2 and *Benefit-of-Faster-Development-%* = 3, the total number of zygotes produced will be: 100 – 2 + 3 = 101, where the decrease (2) is randomly applied to the whole offspring, whereas the increase (3) is randomly applied to one of the three genotypes of the offspring (i.e., either to *++*, *+e*, or *ee* zygotes). As a second example, consider a plate low in bacteria but containing *C. elegans* larvae, and a cross between two EU worms, a *++* and a *+O*. If *Cost-of- Starvation-%* = 1, *Cost-of-Plasticity-%* = 3, *Benefit-of-Predation-%* = 2, and if the first worm eats 2 prey and the second one eats 1 prey, the total number of zygotes produced by the hermaphrodite will be: 100 – 1*2 – 3*2 + 2*(2 + 1) = 98, as both costs and the benefit are applied based on the condition of both parents. More specifically, the total decrease (8) is applied in a completely random way to the entire offspring, whereas the total increase (6) is randomly applied to one of the three genotypes of the offspring (i.e., either to *++* or *+O* zygotes).

The production of the correct number of zygotes and the application of the correct genotype ratios were confirmed by preliminary runs including parents with alternative genotypes, a variety of parameter values for all costs and benefits, and both dietary regimes, i.e., abundant bacteria and no larvae, and low bacteria and presence of larvae. This analysis was conducted using a single *P. pacificus* hermaphrodite for each run (selfing), a single hermaphrodite and a single male (cross), or removing all individuals from the simulation grid except one or two, to test subsequent generations after the first one. The number of offspring and their genetic composition were monitored using numerical monitors and line plots in the interface, representing population size and genotypes.

##### 3.3.9 Custom reporters

Reporters are NetLogo terms which are used to report values. Special procedures can be created to define new, custom reporters that can be called within other procedures. Table S4 contains a list of all custom reporters used in the model.

**Table S4.** Custom reporters defined in the model.

| **Reporter** | **Description** |
| --- | --- |
| *Probability-of-developing-EU-++* | Probability to develop the EU phenotype for *++* nematodes. It reports either 80 or 80 + *Degree-of-Plasticity-%* depending on the abundance of bacteria. Nematodes which do not develop the EU phenotype become ST |
| *Probability-of-developing-EU-+e* | Probability to develop the EU phenotype for *+e* nematodes. It reports either 25 or 25 + *Degree-of-Plasticity-%* depending on the abundance of bacteria. Nematodes which do not develop the EU phenotype become ST |
| *Probability-of-developing-EU-+O* | Probability to develop the EU phenotype for *+O* nematodes. It reports either 25 or 25 + *Degree-of-Plasticity-%* depending on the abundance of bacteria. Nematodes which do not develop the EU phenotype become ST |
| *Generations* | Generations are non-overlapping. The generation counter is updated by 1 unit every 820 ticks (i.e., 82 hours) |
| *Population-size* | Reports the total number of *P. pacificus* nematodes during simulations |
| *Hermaphrodites* | Reports the number of hermaphrodite (XX) *P. pacificus* nematodes |
| *Males* | Reports the number of male (XO) *P. pacificus* nematodes |
| *Frequency-of-++* | Reports the relative frequency of *P. pacificus* nematodes with genotype *++* |
| *Frequency-of-+e* | Reports the relative frequency of *P. pacificus* nematodes with genotype *+e* |
| *Frequency-of-ee* | Reports the relative frequency of *P. pacificus* nematodes with genotype *ee* |
| *Frequency-of-+O* | Reports the relative frequency of *P. pacificus* nematodes with genotype *+O* |
| *Frequency-of-eO* | Reports the relative frequency of *P. pacificus* nematodes with genotype *eO* |
| *Frequency-of-+* | Reports the relative frequency of the *+* allele |
| *Frequency-of-e* | Reports the relative frequency of the *e* allele |
| *++* | Reports the number of nematodes with genotype *++* |
| *+e* | Reports the number of nematodes with genotype *+e* |
| *ee* | Reports the number of nematodes with genotype *ee* |
