## Supplementary File 2 for "Agent-based modelling of a nematode system provides novel insights into the evolutionary constraints and modulators of phenotypic plasticity, bet-hedging, and environmental homeostasis"

### Relevant details about *Pristionchus pacificus* for the implementation of the agent-based model

*P. pacificus* nematodes are the main agents of *PhePlastiComp*. After its discovery in Pasadena, CA in 1988 and its description as a novel species some years later (Sommer et al., 1996), this animal has been used as a comparative species to *C. elegans*, and its genomic information has been considered an important element to fill the gap between *C. elegans* and the diversity of nematodes. In other words, it has been conceived as a satellite species (Hong & Sommer, 2006; Sommer R. , 2006; Srinivasan & Sternberg, 2008; Schroeder, 2021). However, in more recent years, *P. pacificus* has also become an established, successful model organism in evolutionary biology, especially for the investigation of phenotypic plasticity (Gilarte et al., 2015; Dardiry et al., 2023). I summarise below some of the general characteristics of this species, focusing on those that are relevant to this study. Further details are provided in Supplementary File 1 (ODD protocol).

*P. pacificus* (family: Diplogasteridae) is a cosmopolitan nematode species that lives preferentially on scarab beetles and whose size at the adult stage is about 1 mm (Mitreva et al., 2005; Sommer et al., 2017). It can be easily reared on bacterial cultures in the laboratory (Schroeder, 2021), mainly due to its 4-day life cycle at 20°C (Sommer R. , 2006). Before reaching adulthood, it experiences four different juvenile stages (J1, J2, J3, and J4), each of which ends with a moulting event. J1 larvae moult to J2 before hatching (Hong & Sommer, 2006; Schroeder, 2021). *P. pacificus* worms have an androdioecious mating system, as populations are composed of self-fertilising hermaphrodites (XX) and a small percentage of males (XO), whose rates of production were proven to be temperature-dependent (Pires-daSilva & Sommer, 2004; Morgan et al., 2017). Since selfing is a strong type of inbreeding, the prevalence of hermaphroditic individuals can lead to a clonal population (Avise, 2011), that is a feature of critical importance for experimental aims.

In the context of this work, the most relevant peculiarity of *P. pacificus* is the production of its mouth phenotype, that requires a complex interplay between a multitude of genetic, developmental, and environmental factors. Even when the environmental conditions are fixed (e.g., under standard laboratory conditions with no food deprivation or overcrowding), adults with the same genetic background consistently exhibit two alternative mouth forms, that is, eurystomatous (EU) and stenostomatous (ST), with typical, non-identical, and fluctuating ratios observed between different isolates. This indicates that the EU/ST dimorphism is regulated stochastically (Ragsdale et al., 2013; Susoy & Sommer, 2016; Sommer et al., 2017). Among the main divergences between the two morphs, EU worms have a broad buccal cavity, a claw-like dorsal tooth and an opposing sub-ventral tooth, whereas ST worms have a narrow buccal cavity and a single, flint-like dorsal tooth. These differences in morphology were shown to lead to alternative dietary behaviours, with EU worms being facultative predators and ST worms exclusively microbial feeders (Sommer R. , 2006; Serobyan et al., 2013; Sommer et al., 2017; Sieriebriennikov et al., 2018).

The dimorphism is related to a binary, irreversible developmental decision made at the final moult from J4 to adulthood and induced by a gene acting as a switch, the sulfatase-encoding *eud-1* gene, that directs a gene regulatory network consisting of a set of proteins which affect mouth formation. Interestingly, the expression of this switch is regulated by several environmental factors and epigenetic mechanisms, that may cause the EU/ST ratio to deviate from its typical values (Dardiry et al., 2023). So, this species shows a fascinating combination of both stochastic and conditional regulation of phenotypic plasticity (Sommer et al., 2017). In particular, *P. pacificus* mouth dimorphism is a polyphenism, that is, a kind of plasticity where organisms show discrete, distinct developmental outcomes rather than a continuous range of responses along an environmental gradient (Bento et al., 2010; Fusco & Minelli, 2010; Yang & Pospisilik, 2019).

Environmental variables regulating mouth polyphenism include nutrient availability, temperature, and adult crowding (Bento et al., 2010; Werner et al., 2018; Lenuzzi et al., 2021). It was also shown that both mouth phenotypes are adaptive, each one under specific surrounding conditions: ST nematodes develop faster, particularly in contexts with abundant microbial food, whereas EU nematodes can exploit alternative food sources (e.g., nematodes of other species) when bacteria are scarce (Serobyan et al., 2014; Susoy & Sommer, 2016). Finally, it is now well-acknowledged that both mouth-form ratio and environmentally-induced changes in this ratio can be altered by genetic changes. In particular, some dominant, loss-of-function mutations affecting the *eud-1* gene were shown to determine a constitutive expression of the ST phenotype in the lab, whereas additions of *eud-1* copies or overexpression of *eud-1* resulted in the exclusive expression of the EU phenotype (Ragsdale et al., 2013; Susoy & Sommer, 2016; Sommer et al., 2017; Sieriebriennikov et al., 2018). This means that, in principle, it is possible to conceive experimental scenarios involving both plastic and non-plastic (or dimorphic and monomorphic) strains/genotypes of *P. pacificus*, depending on the state of *eud-1*, that is a focus of *PhePlastiComp*.

Indeed, differently from previous research (Kalirad & Sommer, 2024), in this study non-plastic strains are produced by mutations affecting the *eud-1* gene. However, I do not consider known mutations disrupting the polyphenism: in the model, non-plastic strains include mutants resulting from any hypothetical mutation that affects *eud-1* and has the exclusive effect of suppressing mouth dimorphism (which can only be observed in plastic strains (Casasa et al., 2023)). This choice was made to keep the generality of the model broader and because laboratory mutants with induced *eud-1* mutations might show reduced fitness when compared to wild-type worms. Future field studies could support the presence of strains carrying *eud-1* mutations with no negative effects on fitness, that could eventually provide further empirical data for comparisons with the outcomes of my simulations.
