## Supplementary File 3 for "Agent-based modelling of a nematode system provides novel insights into the evolutionary constraints and modulators of phenotypic plasticity, bet-hedging, and environmental homeostasis"

Macro-validation and preliminary runs

An expected, straightforward consequence of the overwhelming prevalence of selfing as a mode of reproduction is the abrupt collapse of the heterozygous genotype in a short time, leading to the exclusive presence of homozygous pure lines (Guzella et al., 2018; Cornetti et al., 2021). The scenario “Decay of heterozygosity” informs that, using a population of 300 nematodes (initial composition: 100 *++*, 100 *+e*, and 100 *ee*) within a context devoid of costs, benefits, environmental variability, and non-disjunction events, and with *Carrying-Capacity* = 300, heterozygotes disappear very quickly from the simulation lattice in all runs: the average time to disappearance was 7.42 generations (σ = 1.88).

To test whether the two parameters *Benefit-of-Faster-Development-%* and *Benefit-of-Predation-%* were in line with empirical evidence suggesting adaptive advantages associated with faster sexual maturation of ST worms and with the ability of EU worms to exploit alternative food sources (Serobyan et al., 2014; Sommer et al., 2017), respectively, I used the scenario “Neutral condition” as a baseline. Considering an initial symmetric condition (i.e., 150 *++* and 150 *ee*, with *Carrying-Capacity* = 300) and excluding costs, benefits, environmental variability, and non-disjunction, an approximately equal frequency of fixation of both alleles was observed, with a relative frequency of fixation of *+* (henceforth, *ƒ_fix_(+)*) = 52%, and *ƒ_fix_(e)* = 48%, which confirms the selective neutrality of the alleles. Keeping in mind the mouth-form ratios associated with each genotype (see Table 1 and Supplementary File 1), benefits in fecundity favouring the ST phenotype are expected to increase the frequency of *e*, whereas benefits favouring the EU phenotype will mainly increase the frequency of *+*.

The scenario “Effects of benefits to ST worms” differs from the “Neutral condition” only for the introduction of a gradually increasing benefit in terms of fecundity assigned to ST nematodes (1%, 2%, and 3%). Each of these benefits is associated with a set of 100 runs. In this case, *C. elegans* larvae are excluded from the simulation grid. The runs show that *Benefit-of-Faster-Development-%* can increase the frequency of fixation of *e*, while decreasing its average time to fixation (1%: *ƒ_fix_(e)* = 97%, mean time to fixation of *e* (henceforth, *t̅_fix_(e)*) = 280.32, with IQR = 217, Q1 = 154, Q2 = 240, Q3 = 371; 2%: *ƒ_fix_(e)* = 98%, *t̅_fix_(e)* = 209.81, with IQR = 128.75, Q1 = 130, Q2 = 171.5, Q3 = 258.75; 3%: *ƒ_fix_(e)* = 100%, *t̅_fix_(e)* = 148.6, with IQR = 84.25, Q1 = 96.5, Q2 = 128.5, Q3 = 180.75).

The scenario “Effects of benefits to EU worms” is also based on the “Neutral condition” above, this time considering a gradual increase of values of the parameter *Benefit-of-Predation-%* (1%, 2%, and 3%, 100 runs for each value). This benefit is only experienced by EU nematodes due to predation of *C. elegans* larvae which, in this case, are present in the system, and is associated with a limited ability to kill each prey (*Efficiency-of-Attack-%* = 8%). This probability of killing prey was chosen because it led to outcomes that are comparable to those obtained in the previous scenario. This scenario seems to support the general tendency that the higher the increase in fecundity for EU worms, the greater the number of fixations of *+*, with decreasing average time to fixation of *+* (1%: *ƒ_fix_(+)* = 94%, *t̅_fix_(+)* = 259.57, with IQR = 175.25, Q1 = 146, Q2 = 210, Q3 = 321.25; 2%: *ƒ_fix_(+)* = 99%, *t̅_fix_(+)* = 192.33, with IQR = 127, Q1 = 111.5, Q2 = 176, Q3 = 238.5; 3%: *ƒ_fix_(+)* = 99%, *t̅_fix_(+)* = 142.87, with IQR = 67.5, Q1 = 98, Q2 = 132, Q3 = 165.5).

Finally, in the scenario named “Effects of intrinsic costs of plasticity”, I tested whether increasing costs of plasticity (1%, 2%, and 3%) had a penalising impact on the fixation of *+*, that is an expected implication within a system where the ability to produce and maintain plastic responses affects fitness in a negative way (Dardiry et al., 2023). This impact was observed in all three experimental sets (1%: *ƒ_fix_(+)* = 3%, *t̅_fix_(+)* = 143.33; 2%: *ƒ_fix_(+)* = 1%, *t̅_fix_(+)* = 82; 3%: *ƒ_fix_(+)* = 0%. Except for the variation of the parameter *Cost-of-Plasticity-%*, this scenario is also identical to “Neutral condition”.

Collectively, the behaviour of the modelled system reflects, at a qualitative level, general patterns which would be expected in a real population sharing the same basic characteristics.
